# A parsimonious murburn model for microbial motility connects metabolic water ejection to observable mechanical outcomes

**DOI:** 10.64898/2026.06.06.730539

**Authors:** Kelath Murali Manoj, Abhijith Anandakrishnan, S. Sunil Kumar, Daniel Andrew Gideon

**Author notes:** Corresponding authors: Kelath Murali Manoj; Abhijith Anandakrishnan.

## Abstract

The classical model of bacterial flagellar motility posits a rotary engine driven by proton motive force (pmf), with torque generated by stator–rotor interactions and transmitted through a flexible hook to a helical filament. Despite decades of acceptance, this model faces fundamental challenges in thermodynamics, structural mechanics, evolutionary parsimony, and direct observational evidence. We develop and quantitatively test the murburn model, a new paradigm for bacterial motility in which water, produced as an inevitable byproduct of metabolic redox activity, is ejected via the basal secretory module and channelled along the spiral grooves of the flagellar filament. The ejected flow creates a local shear field that induces a transverse bending wave; the precession of this wave is observed as apparent rotation and generates thrust through anisotropic viscous drag, without any rotary motor, ion gradient, or axial rotation. The principal contribution of this work is a self-contained, first-principles treatment of this mechanism: for a unipolar flagellated cell we derive the governing low-Reynolds-number elastohydrodynamic relations from slender-body theory and show that physiologically realistic rates of metabolic water production reproduce the observed swimming speeds and apparent-rotation frequencies at a small fraction of the cellular energy budget, while direct jet propulsion is quantitatively excluded. Building on this derivation, we provide a force-balance comparison of the competing propulsion mechanisms, obtain a set of falsifiable predictions that distinguish the murburn model from the rotary motor, and report a structural analysis of cryo-EM flagellar-hook architectures that reveals solvent-accessible radial canals consistent with lateral water transport. The same single principle accounts for swimming, tumbling, gliding, spirochete undulation, and archaeal motility, without requiring rotating shafts, ion-gradient coupling, or complex switching mechanisms.

## I. Introduction: The classical rotary model and its entrenched status

We shall call the basal module and filament of any bacteria as the bacterial flagellar system (BFS). When referring to the classical rotary perception, we shall call it as bacterial flagellar motor (BFM).

### I.1 What is currently believed

The classical model solicits mechanical rotation (needs gaps for the rod to pierce through several phases), membrane integrity (needs these piercings to be sealed), and simultaneous ion-gradient generation and utilization (needs phase-isolation and proton-cycling). It is believed that respiratory proteins pump cytoplasmic protons into the periplasm and the very protons return to the cytoplasm via the plasma-membrane embedded Mot-AB ‘stator’ units, twirling the C-ring of the BFS, which rotates the whole length of the flagellum along its longitudinal axis. The various necessities for the successful execution of this proposal are in conflict with each other and do not find justifiable parallels in any naturally self-assembled or manmade synthetic systems. The salient aspects of the prevailing model (Berg, 2003; Berg and Anderson, 1973; Wadhwa and Berg, 2022) rests on the following claims, as shown in Table 1.

**Table 1.** The salient perceptions for motility function within the classical bacterial flagellar motor (BFM) model.

| Component | Function | Structural rationale |
| --- | --- | --- |
| Respiratory proteins<br>(Non-BFM components) | Pump and cycle protons for powering the BFM | Trans-membrane unidirectional proton-conduits |
| Tapping of proton motive force (*pmf*) by BFM | Proton flow drives rotation | MotAB conserved mechanism |
| Membranes, Cell wall, MS, L & P rings | Sealed firmaments and ‘bushes’ | Snug-fitting proteins with radial symmetry/assembly |
| C-ring (FliG/FliM/FliN) | 1 Rotor per basal module; rotates relative to stator | Variable subunits of the three proteins, ~40 to 65 nm in diameter |
| MotA/MotB complex | Multiple stators; anchored to peptidoglycan, conducts protons or cations | 5 MotA + 2 MotB per unit; 11–18 units per motor |
| Rod, hook, filament | Torque transmission from rotor to propeller | Hollow, flexible, polymorphic |
| Direction switching | CheY-P binding to FliM serves as clutch | Reverses from CCW to CW by altering motor-stator interaction |

### I.2 The core logical problem

The classical model requires that a self-assembled supramolecular complex spanning 5–7 distinct phases (cytoplasm, inner membrane, periplasm, peptidoglycan, outer membrane, extracellular space) achieves continuous, high-speed rotation without leakage, with perfect seals at sub-Angstrom scales, while simultaneously exporting flagellin monomers through the same hollow channel (Evans et al., 2014; Macnab, 1999, 2003). (It must be noted that the hollow connectivity of the filament connects the inside and outside phases and this is not seen to perturb pmf or mechanics of the overall BFM model.) No man-made or naturally occurring self-assembled system demonstrates such a combination of features; as protons (on which a vast majority of these purported protein-assemblies or motors are supposed to run!) are less than 2 femtometres in dimension and atoms are of minimally about 2 Angstroms in dimension. The hydrated ions of proton (are a statistical distribution of various levels of hydrations; like Eigen and Zundel-averaging about 50% and 30%, respectively) and sodium are much larger. These decades-old spectroscopically well-established facts of physical chemistry do not find any consideration in the classical biology models; and the Grotthuss mechanism is misinterpreted to conveniently (erroneously!) serve as a pan-systemic mechanistic rationale. (That is-it is projected that a proton is present in one phase for gradient-generation but absent for gradient-dissipation, at the same instant, for proton-cycling!) Also, the same basal module structures are supposed to work efficiently, both with protons and sodium ions, which have vastly different thermodynamic (redox and hydrational), dimensional and mobility properties (Asai et al., 1999). At one hand, the classical model seeks selectivity/specificity and it also espouses degeneracy at the other.

### I.3 The proton-centric Mitchellian bioenergetic postulates have been debunked

The rotary model is deeply rooted in Mitchell s chemiosmotic theory (Mitchell, 1979), which claims that proton pumping by electron transport chains (ETC) creates a transmembrane pH gradient and electrical potential (*pmf*) that drives Complex V’s rotary ATP synthesis and flagellar rotational motility. However, as demonstrated extensively in our previous works (Manoj, 2018a,b,c, 2020a,b,c; Manoj and Bazhin, 2021; Manoj and Gideon, 2022; Manoj et al., 2019a,b, 2020, 2021, 2022a,b, 2023a,b,c,d), the Mitchellian framework is thermodynamically, kinetically, mechanistically, structurally and evolutionarily untenable.

Key arguments include:

#### A. Proton availability

At a given instant, the physiological pH (7.5) affords in a 0.4 femtoliter bacterial cell approximately *<*10^1^ free protons total (Manoj et al., 2023a). The classical model purportedly requires ~1200 protons per rotation (Milo and Phillips, 2015) of the C-ring of one flagellum; an impossibility from an instantaneous perspective.

#### B. Absence of pH gradient

Most bacteria maintain cytoplasmic pH 7.5–7.7 with periplasmic pH also near neutrality (Wilks and Slonczewski, 2007), and the latter phase is equilibrated with the external medium. Alkaliphilic bacteria swim and thrive at external pH 11–12 with cytoplasmic pH 7-8, directly contradicting *pmf* logic (Aono et al., 1992; Krulwich, 1986).

#### C. Misinterpretation of uncoupling data

The BFS research took a major detour with the interpretation of data with “uncouplers”. Comprehensive analyses of data shows that molecules like DNP and CCCP do not act as protonophores but as DRS modulators/scavengers, as their uncoupling activity does not correlate with pK_a_ or protonability (Manoj, 2018b; Manoj et al., 2023b; Terada, 1990).

In this work we move beyond critique to develop and quantitatively test a positive alternative; our specific, original contributions are: (i) a first-principles, low-Reynolds-number elastohydrodynamic model in which metabolically generated water, ejected at the basal module and guided by the filament’s spiral grooves, drives a precessing bending wave that an observer reads as apparent rotation (Section III); (ii) a closed quantitative chain showing that physiologically realistic water-ejection rates reproduce the measured swimming speeds and apparent-rotation frequencies at a small fraction of the metabolic budget, together with a force-balance analysis that quantitatively excludes direct jet propulsion (Sections III.8–III.9); (iii) a set of falsifiable predictions that distinguish the murburn model from the rotary motor (Section IV); and (iv) a re-examination of publicly deposited cryo-EM structures indicating solvent-accessible inter-subunit pores consistent with lateral water transport (Section IV.1).

## II. Elaborate foundational criticisms of the rotary BFM model

### II.1 Thermodynamic impossibility highlighting the proton cycling

Let’s accede to the classical BFM requisites and say that the bacterial cytoplasm is not practically aprotic and assess the dynamics of proton cycling. Based on gearing principles and the structural stoichiometry of the bacterial flagellar motor, if the FliM-containing C-ring rotates at ~2 × 10^3^ Hz, the MotA assembly of the MotA-B protein complex (the stator) must rotate at approximately 13,600 Hz (or ~1.3 × 10^4^ Hz). This is derived from taking a gear ratio of 6.8 to 1 between the stator and the rotor. [[Determining the gear ratio in the BFS: The rotation of the ‘flagellar motor’ is understood to result from the function of a set of intermeshed gears where the smaller “drive gear” (the MotA-pentamer rotating against a static MotB-dimer; both together called the stator) turns the larger “driven gear” (the FliG/M C-ring). The rotor (C-ring): Recent high-resolution cryo-EM studies show that the C-ring typically consists of 34 subunits of FliG/FliM. So, 34/5 = 6.8]]

So, the number of protons flowing in per second through a single BFS is: (1.3 × 10^4^) rotations of a single MotA-B x (2 × 5 protons) per rotation of MotA-B) x 12 MotA-B units per C-ring = *>*10^6^ protons per second at a single C ring. There could be even 10 flagella per cell, if peritrichous. This dictates that the single bacterium should be able to cycle *>*10 million protons across its membrane per second. The maximal conceivable respiratory proteins (Complexes I, III and IV) range in the order of 10^3^ to 10^4^ per bacterial membrane. Even if there is a selective ion pump that could pump protons (which is infeasible, as protons are too small to afford any ‘select-ability’!), Complex V would partake a significant portion (let’s say, at least 1/10^th^, by any conservative estimate) of these protons for ATP-synthesis (which would be needed for routine cellular work and maintenance). It is known that protons require milliseconds to cross membranes. This would mean that the number of protons dynamically required (*>*10^7^) for cycling the cell through its ATP-synthesizing cum motility agenda falls short by orders of magnitude (10^3^ x 10^−1^ x 10^3^ ≈10^5^ per second) that could potentially be provided by any purported pumping.

Once again, from the static perspective, as the bacterial interior is practically aprotic, i.e., it would realistically have ~10^0^ protons. [[(Crowding corrected volume = 10^−16^ L) x (concentration of H^+^ = 10^−8^ moles/L) x (Avogadro’s number = 6 × 10^23^ per mole) = *<*1]]. Even allowing for proton recycling via water auto-dissociation, the oft-quoted Le Chatelier argument fails because the equilibrium constant for H_2_O dissociation (*K* _eq_ = 10^−14^) gives Δ*G* ≈+80 kJ/mol at 27^°^C; the forward reaction is highly unfavorable.

Water does not spontaneously provide protons for pumping; this is when the classical BFM 10^6^ – 10^7^ more protons than the cell can instantaneously supply. Besides, most bacteria maintain their internal pH at 7.4–7.8 regardless of the broad external pH range of the medium (Slonczewski et al., 2009). From a dynamic perspective (acceding to the mandates/claims of the BFM model), the proton requirement still falls at least two orders short of the requirement! Therefore, the proton-based powering/mechanistic logic was/is a thermodynamically unattainable premise.

### II.2 Structural and mechanical implausibility

This is perhaps the most important part, which is evident for all to see, understand and infer. Before anything else, we present the salient features of the relevant structures in the BFS, from Figures 1 through 8, with analytical discussions in sessions that follow.

**Figure 1.**
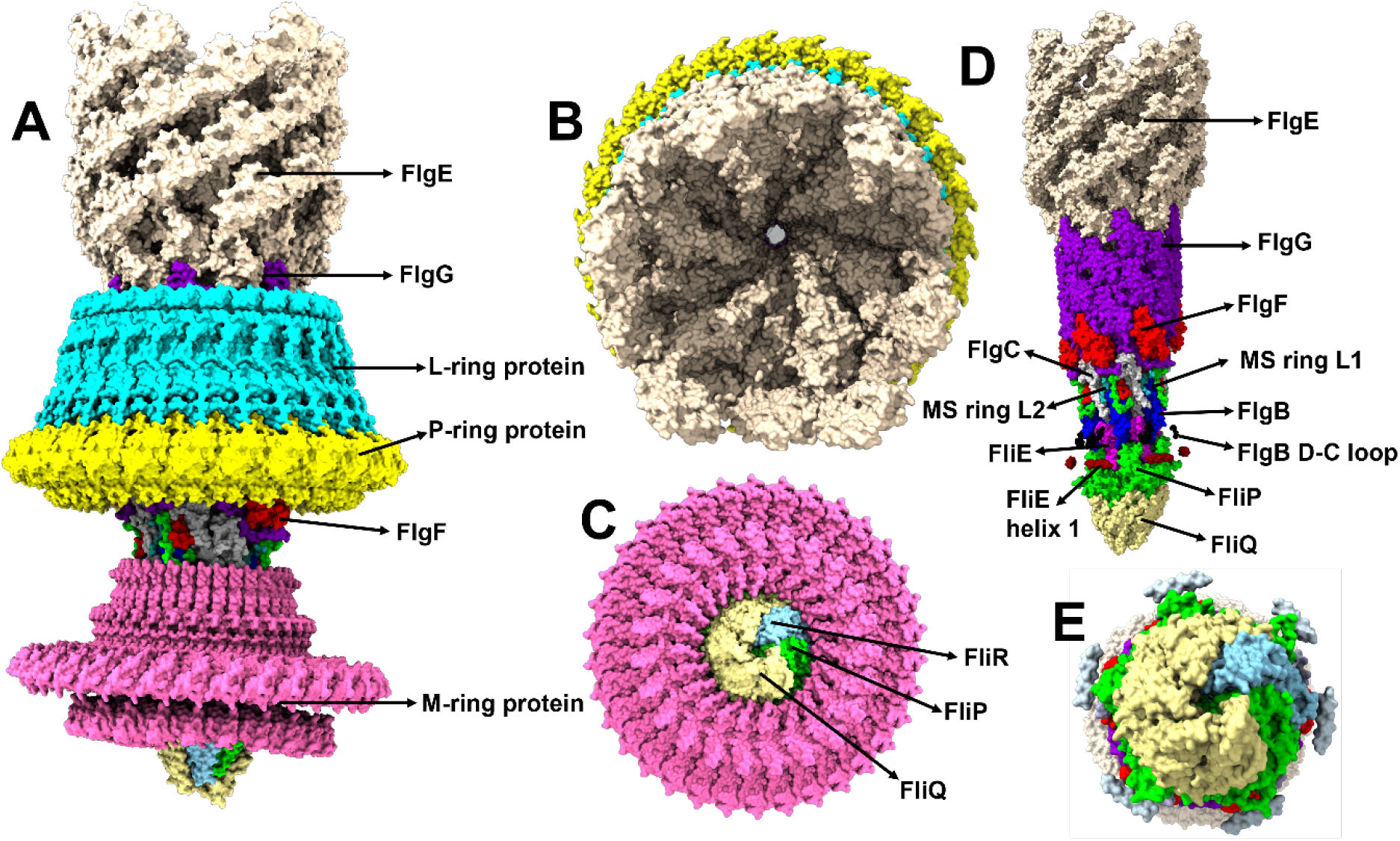
The external components of hook and filament of the BFS. **A –** Overall structure of the Salmonella enterica subsp. enterica serovar Typhimurium str. LT2 “motor hook” complex (PBD ID: 7CGO). The major parts of the complex are labelled and the parts enclosed within the LP and M rings are shown in panel **D**. Panel **B** shows the FlgE hook protein complex and the aperture in the middle appears to be like a tunnel which leads to the tip of the flagellar FliR/FliP/FliQ complex. Panel **C** shows the M ring along with the FliR/FliP/FliQ complex (this complex is magnified in panel **E**).

#### A. The multi-layer rotating shaft problem

The classical model requires a highly coordinated, multi-layered alignments allowing rotations with low-friction, torque-transmitting (10^3^ to 10^4^ pN.nm) assembly spanning heterogeneous phases, which raises legitimate questions about its spontaneous assembly and robustness. Figure 1 and Figure 2 show the external components of the hook and filament of the Salmonella BFS such as LP ring and MS ring. The rod must rotate while piercing the inner membrane (MS ring), peptidoglycan (P ring), and outer membrane (L ring) in Gram-negative bacteria cannot act like bearings and braces at the same time. Also, such a rotation requires:

**Figure 2.**
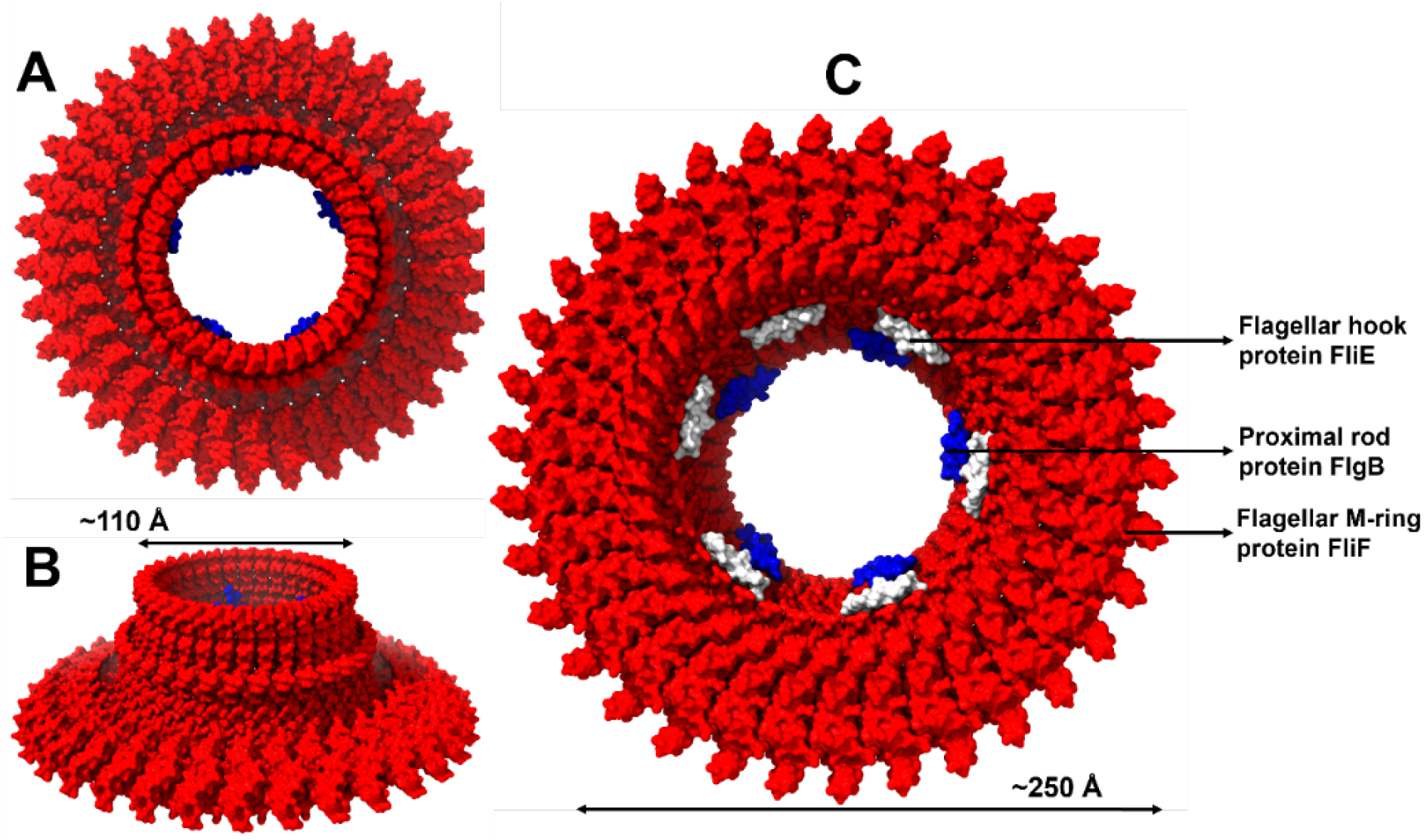
MS ring of Salmonella enterica subsp. enterica serovar Typhimurium str. LT2 (PDB ID: 8WK4). **A & B**. Top to bottom and side views of the complex. **C**. Funnel shaped views of the MS ring, showing FliE (white shading), FlgB (blue shading) and FliF (red).

- Sub-Angstrom (femtometer-level!) precision seals against proton leakage
- Low-friction rotation across hydrophobic and hydrophilic interfaces
- Simultaneous torque transmission through a flexible hook

No known bearing system achieves this at the 2–20 nm scale. Thermal motion (*k* T≈ 4.1× 10^−21^ J at 300 K) ensures gaps exist; fluids leak through nanopores. The classical model s assumption of “bushings” (L-P rings) has no mechanical analog in self-assembled systems (Figure 1). For a detailed discussion on the various facets considerable for rotation in natural or human-assembled systems, please refer to our earlier work (Manoj et al., 2023b).

#### B. The stator rotation speed impossibility

Based on gearing principles and structural stoichiometry (rotor: 34 subunits; stator: 5 subunits; gear ratio = 6.8), if the C-ring rotates at ~2 × 10^3^ Hz (reported for *Vibrio*), the MotA/B stator ring must rotate at- 2000× 6.8 = 13,600 Hz≈ 816,000 or ~ 10^6^ RPM!! Further, a cup of 50 nm across FliM-N rotor (C-ring) rotating at speeds approaching 2000 hertz would create some localized shear and dissipation (even in the low Re regime of bacterial cytoplasm). Since *v* = *ωr*, 2 × 10^3^ x 3 × 10^−8^ = 6 × 10^−5^ m/s. This means that local surface velocity (near the spinning C-ring) would be ~60 microns per second! Along the same coin, the local surface velocity at the MotA complex would be 1.36 × 10^4^ x 3 × 10^−9^ = 4 × 10^−5^ m/s or 40 microns per second, which is orders higher than the magnitude of the bacterium. (This value is large, even if the flow decays rapidly as an inverse squared function! How could the Mot B dimer, which is non-covalently tethered to the peptidoglycan layer above the plasma membrane, and known to be dynamically replaced, hold on to serve as a firmament or true stator, to afford relative rotation for Mot A, which should in turn churn a massive C-ring cum rod assembly against insurmountable mass load and hydrodynamic drag?) A naturally assembled membrane-embedded protein complex rotating at nearly 1 million RPM in a viscous lipid bilayer is incredible and thermodynamically absurd. The frictional dissipation would denature the protein complexes instantly. All such speculations perpetuate only because of the unchallenged perception that BFS is a physiological rotary functionality. Besides, we must ask how torque generation is dependent on directional ion-coupled conformational changes involving the same stator machinery to generate equal torque in opposite directions devoid of any mechanical reconfiguration. The MotAB complex of *C. jejuni* is shown in Figure 3.

**Figure 3.**
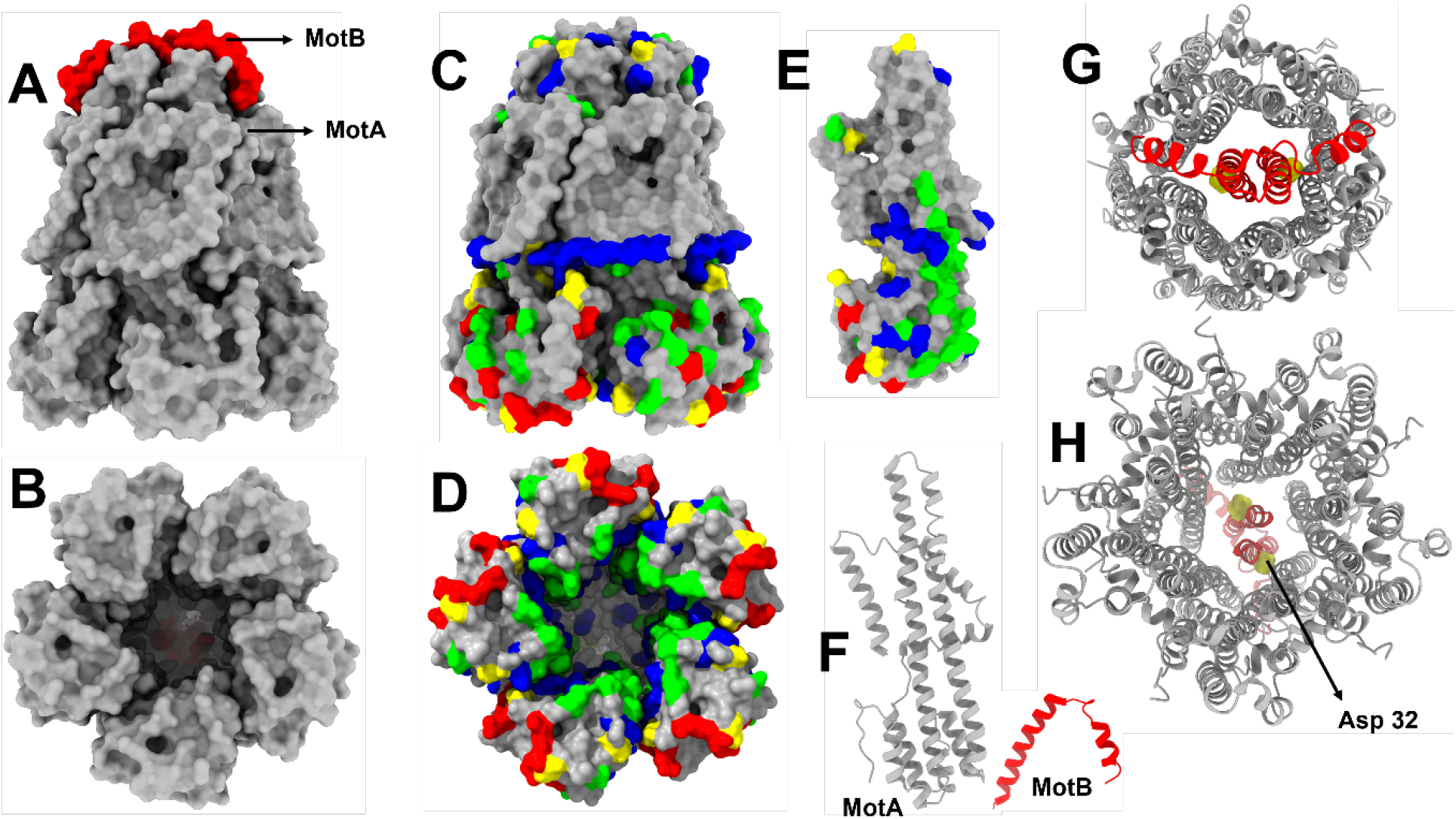
Structure of the C. jejuni MotAB complex (PDB ID: 6YKM). **A** – Overall top-bottom view of the MotAB stator complex, showing Mot A (gray) and Mot B (red) proteins. **B –** bottom view. **C –** MotAB with colored charged residues (Arg-red, Lys-blue, Asp-yellow and Glu-green). These color codes are retained in panels **C-E**. Panel **D** shows the bottom of the complex with different charged residues. The interior of the MotA pentameric complex is lined with both Glu and Lys residues. The bottom is rich in various charged amino acid residues. Panel **E** is the surface view of a MotA monomer. Panel **F** shows the ribbon structures of MotA and MotB. Panels **G** and **H** show the view of the MotAB complex from top to bottom and bottom to top, with the key Asp 22 in this PDB (Asp 32 in literature) coloured yellow.

#### C. The variable stoichiometry problem, an evolutionary imbroglio

If the flagellar motor were a precision nano-machine, the number of stator units and rotor symmetry would be strictly conserved. However, from Table 2, it can be seen that the spectral spread of the proteins across the bacterial population (the actual length of amino acids and their conservation at a definite locus, both!) is quite otherwise! It is inconceivable how the various proteins would effectively coordinate freely reversible movements and clutch-gear roles with such a diversified evolutionary spread. We had also shown analyses of the FliG protein and argued that it is structurally unlikely to serve as the clutch-switch for MotA-Cring interaction, resulting in the CCW to CW rotation-directional change (Gideon et al., 2024).

**Table 2.** A snapshot of some key proteins’ length and copy numbers in the BFS.

| Species | FliN length | FliM length | FliG length | MotB length | Stator numbers |
| --- | --- | --- | --- | --- | --- |
| *E. coli* | 137 | 334 | 331 | 308 | 11 |
| *H. pylori* | 79 | 354 | 343 | 257 | 18 |
| *A. fabrum* | 179 | 321 | 375 | 230 | variable |
| *S. meliloti* | 198 | 315 | 345 | 292 | 11-18 |
| *B. aphidicola* | 134 | 334 | 320 | N/A | N/A |

The chain lengths vary by *>*200% and copy number of key proteins per module varies by 64%. Rotor symmetry varies from 32-36. Stator numbers vary from 11–18. As argued in Gideon et al. (2024)(Section 3.2): “With such diversities in composition and very clear differences in assembly/gross structure, it is clear that a common mechanism, such as H^+^/Na^+^-translocation mediated rotary propulsion/flagellar movement appears highly unlikely.” Most importantly, it is indeed very difficult to envisage that a highly sophisticated motility system would evolve with about 50 gene products (giving as many different proteins), assembling with various ratios (300-400 proteins in various multimeric compositions/numbers and *>*20,000 flagellin monomers; the basal modular assembly weighing around 11 MDa and the filament per se being *>*1 GDa; a total weight of the BFS being more than a billion Daltons, which amounts to1/100 - 1/1000 of the bacterial body weight!), working across more than half a dozen distinct phases, only with *<* 2% efficiency (Purcell, 1997). This is why Michael Behe justifiably called the BFM model as an example of “irreducibly complex” system (Behe, 2004), explainable only with intelligent design (and not spontaneous molecular evolution!). If we say that it is not a complex nanomotor but a simple fluid secretory/ejection system common to diverse life forms (which happens to work efficiently at the nanoscale and low Re realms too!), we can still have an evolutionarily grounded explanation for bacterial flagella-assisted motility.

#### D. The abuse of mechanical terminologies/concepts and the screw paradox

In analogy with rotary electromagnetic motors/generators and mechanical turbines/propellors, the classical BFM models uses rotary drive machining/working terminologies like rotor, stator, bearings, bushings, housings, driveshaft, clutch, gear, torque, ratchet, power-stroke, helical or corkscrew propellor, RPM, etc. Firstly, we present a two-layered (at two hierarchical levels) critique of the highly popular cork-screw propeller analogy-drawing when the bacterial body-geometry is formalized for a polar monotrichous bacterium like *Vibrio cholerae*. From screw theory kinematics, for a left-handed screw/helix (LHS), CCW rotation leads to motion toward base (pulling) and CW rotation affords motion away from base (pushing). Therefore, for a monotrichous flagellum, a CCW rotation would lead to a pulling, not pushing! This violates the basic observational inference in bacterial systems wherein the CCW movement of the polar LHS flagellum apparently propels/pushes the bacteria forward! (We know this for a fact from daily experience that for a macroscopic right-handed screw, CW rotation drives the screw into the material (pushing). Also, for a left-handed screw, CCW rotation pulls it out from the material.) Therefore,

##### Layer 1 critique

Internal inconsistency in the classical BFM narrative! The classical pedagogic explanation often says that the flagellum acts like a corkscrew/propeller. Under that analogy, LHS + CCW should make the bacterium exhibit pulling of the bulk body, quite unlike the pushing that is naturally observed. So, the classical explanation fails on its own stated intuitive mechanical basis. This is a rhetorical/internal critique.

##### Layer 2 critique

Even the screw analogy itself is incomplete at low Re! At bacterial scales, inertia negligible, and it is acknowledged that propulsion arises from anisotropic viscous drag, waveform propagation matters. Therefore, from a hydrodynamic perspective, macroscopic screw intuition is insufficient anyway.

The model mixes macroscopic intuition with micro-hydrodynamics in an inconsistent manner. We would like to add the disclaimer here that we do not question the theoretical premises that at low Reynolds number regimes, there is little inertia and motion would critically depend on non-reciprocal (and anchoring) forces, anisotropic viscous drag, elastic constraints and waveform propagation. It is just that the classical model requires more/hidden assumptions to reconcile helix handedness and thrust direction. Secondly, for a peritrichous organism like *E. coli*, from geometry (strictly), LHS flagella rotating CCW located at different positions/orientations would produce force vectors along its own axis, which are not naturally aligned. Furthermore, these flagella do not have any resemblance to helical cork-screw type structures! We do concede that at low Re, long helices can interact via viscous forces and can self-organize into bundles owing to hydrodynamic instability. But the concern is that motors are independent and lack coordination. Phase-locking would require precise torque balance, elastic compliance and fluid coupling, which is not trivial or guaranteed. Particularly, flagella on two opposing sides of the bacterium would generate non-productive divergent force vectors, whose synchronization is not guaranteed. Therefore, a lot more is desired for the functional justification of the cork-screw analogy in such systems. Thirdly, archaeal flagella, also known as archaella, are fundamentally different from bacterial flagella. They are thinner and assembly-wise more similar to bacterial Type IV pili, lacking the hook-filament-motor structure found in bacteria. Also, some bacteria move across surfaces without flagella at all, using a process called gliding, which involves specialized secretory systems and surface filaments. The classical rotary BFM model needs to resort to multiple phenomenological explanations to address each of these instances. Finally, we address the exact mechanism which is supposed to onset/initiate the rotary movement within the primary component (which is located paradoxically within the so-called stator, the MotA pentamer!). A proton is supposed to trip a conserved acidic aspartate (D32) residue within the MotAB unit. Two relevant questions are – a) Why does the protonation of D32 (a specific aspartate residues) alone occur when there are a myriad protonatable resides present in the entire flagellar assembly (and specifically in MotA/B complex)? b) How does deprotonation occur in time for the next proton to protonate this residue? Can shifts in p*K* _a_ occur quickly enough to restore D32 for the next protonation event? To fact check the claim that D32 is conserved across all bacterial species, we exposed that contrary to such enforced consensus, this was NOT CONSERVED and in fact, the MotAB units had very divergent sequences (see Table 4 of Gideon et al., 2024), some with no aspartate/glutamate for #32±4 residues and some with multiple such residues! So, there is no evident chemical mechanism for chemo-mechanic or ionic “ratcheting” to give the “power-stroke” for the MotA monomer to move 72 degrees with the movement of two protons or sodium ions. Also, there is no real mechanism afforded by BFM advocate till date as to how a protonic (which is actually non-existent!) or ionic or electrical trans-phase gradient could be tapped by a spongy-soft lipid-protein assembly for conversion into rotary motion, particularly when there is no paramagnetic or ferromagnetic sensing/transducing agent within the same. (Quite simply, is the mechanism fundamentally chemical or electrical or is it mechanical? If a combination of all these, how? 5 decades is a long time for any leading researcher to answer such questions.) The earlier consensus in the field was a 4:2 MotA:MotB ratio as this would have made some quantitative/physical sense of a rotary motion from the mechanical movement of two protons/ions. But the later structural finding that MotA:MotB ratio is actually 5:2 across diverse systems clearly does not afford any conceivable/repeatable quantitative/physical mechanism for a 72 degrees ratcheting/power-stroke movement of MotA (from a symmetry perspective). We also argue that quoting cryo-EM images as evidence for MotA subunits’ rotation is misplaced as these images are actually statistically averaged snapshots (and the membrane proteins are squishy-squashy, with very many low-energy domains), and not dynamic photographs! The “motor” metaphor/analogy is a post-hoc conceptual framework, not a demonstrated mechanism. It was imported from engineering (man-made rotary systems) before the molecular structure or mechanism was fully characterized. The consensus narrative was built partly on the metaphor itself. True rotary motors (e.g., turbines) convert energy into continuous, unidirectional torque about a fixed axle. The flagellum s “rotation” is inferred largely from tethered-cell assays and bead rotation experiments; both of which measure whole-cell or filament behavior, not direct molecular rotation at the basal body. This is discussed in the following section that follows, wherein we argue that what is observed is helical wave propagation through a semi-rigid polymer, and the basal body may be the transducer of a conformational relay, but definitely not a rotating machine.

### II.4 The (in)direct evidence for filament rotation: Fact or optical illusion?

#### A. Resolution limits

The flagellum diameter is ~20 nm. The wavelength of light used for optical visualization = 400–700 nm (10–35× larger). Advanced super-resolution microscopy may achieve ~50 nm resolution (Schermelleh et al., 2010), which is still significantly higher than the cross-sectional dimension (the axis around which the rotation purportedly occurs!). As the dimension of the flagella is easily *<*1/10 of the wavelength of light used for visualization (200 to 600 nm), there is no way to ascertain the rotation of the filament through optical means.

#### B. The demonstration by Ali et al. (2016)

In the 69^th^ annual meeting of the APS division of fluid dynamics (Ali et al., 2016), the proceedings are clearly available for all to visually verify. It shows that self-assembled flagellin filaments (no basal motor, no cell body; the auto-assembled are merely tethered at one end to the microscopic glass slide) exhibit apparent rotation/corkscrew motion under external shear flow. In the real world, by all conceivable norms of reality, a tethered filament cannot rotate! Therefore, the observed motion must be a passive hydrodynamic effect of spiraling/revolution owing to fluid shear flows, and not due to axial motor-type rotation sponsored at the basal module (by Cring/MotAB rotation).

#### C. Tethered cell/bead experiments (Berg’s toy mouse analogy)

In the famous video of the pioneer Berg (https://www.youtube.com/watch?v=ioA1yulA-t8), he shows that when a toy mouse’s (whose hands rotate when the body is held by the viewer) hands are held, the body rotates with respect to the held-hands (but this is at the very same frequency, but in opposite direction to what the hands rotate when the body is held!). He argues this proves flagellar rotation. However:

- The tethered cells rotate at 10^0^ to 10^1^ Hz. Physiological flagellar rotation is claimed to be several orders of magnitude higher, at 100 to 2000 Hz. The “higher load” argument that Berg quotes cannot explain this discrepancy in low Reynolds number regimes, where inertial forces are negligible. We should observe significantly higher rotational frequencies in this modality. We reason this observation in a different way. When the filament is held/tethered on to the slide, the resulting shear forces could disrupt normal physiological articulations, releasing the BFS assembly’s snug-fit hold on the filament (implying a loose interaction between the basal module and membrane), and the ensuing fluid dynamics could rotate the body of the bacterium at very small frequencies. This observed reality has little extrapolative merit to propose that bacterial flagella rotate around their longitudinal axes at frequencies of thousands of Hertz (and that this rotation originates owing to a spinning of the Cring at the basal module!), to explain for the high motility of bacteria at low Re.
- The flagellum supposedly switches between CW and CCW rotation on millisecond timescales via the C-ring (FliM/FliN/FliG) responding to CheY-P (Figure 4). Figure 5 shows the fundamental unit of the C-ring, involving interactions between FliG, FliM and FliN. The molecular intelligence of this process (how or why?) is yet unanswered. In rotary motor terms, this means the entire rotor reverses direction nearly instantaneously. No macroscopic rotary motor reverses direction this cleanly. Although the flagellum s moment of inertia is tiny, the structural reorganization implied in the switch is enormous. We argue that the switching mechanism could be more consistent with a two-state functionalism, and not with any change of rotational direction of the basal module, per se. A pertinent question to ask in this regard is, are the non-covalent interactions between FliG in the MS ring (Lys264, Arg281 and Asp289) enough to stabilize binding with MotA’s Arg90, Glu98 (and other charged residues) strong enough to maintain integrity during purported rotation at 10,000–100,000 rpm? How fast must these bonds be made and broken, particularly in the CW and CCW orientations? It is also not clear how transient cooperative binding (similar to oxygen binding to myoglobin) of relatively small signalling proteins like CheY-P contribute such a large and rapidly rotating MS ring complex to trigger a coordinated and almost instantaneous reversal of torque direction without even slightly disrupting motor function. How does CheY-P dissociation reverse the direction of movement once again? The idea that CheY and CheY-P exists in dynamic equilibrium, subject to signalling cues, does not explain how conformational information is propagated around the C-ring. Enough CheY-P dephosphorylation must occur through CheZ dephosphorylation, FliG must rearrange, the rotor-stator interactions should reverse, and all these events must occur while the motor is still spinning at a high speed, requiring a highly cooperative allosteric transition mechanism.
- Torque values (~1000–4600 pN ·nm) are derived from bead-rotation assays and back-calculated using hydrodynamic models of filament drag. These models assume the filament is a rigid helix rotating about a fixed point. The hook-filament junction behaves more like a propagating deformation system. The hook (FlgE polymer) is famously flexible and acts as a universal joint. The hook s supercoiling geometry changes with swimming mode (run vs. tumble), suggesting it s actively participating in shape-change, not passively transmitting torque. But in a true rotary motor, rigidity at the drive shaft would be solicited. Why would evolution retain a flexible joint? The filament-hook system is more consistent with a peristaltic or wave-propagating actuator similar to cilia in eukaryotes before the power-stroke model was settled than a propeller mechanically spun from the base at high RPMs.
- The flagellar filament is a polymorphic helix and it changes its helical parameters under mechanical stress (Calladine, 1976; Hasegawa et al., 1998; Namba and Vonderviszt, 1997). If the filament is deforming while “rotating”, the torque inference is built on an incorrect geometric model. What s measured is angular velocity of a tethered bead, and the torque is a model-dependent back-calculation. The data are also consistent with helical deformation propagation producing net bead rotation without requiring a rotating axle at the base of the BFS.

**Figure 4.**
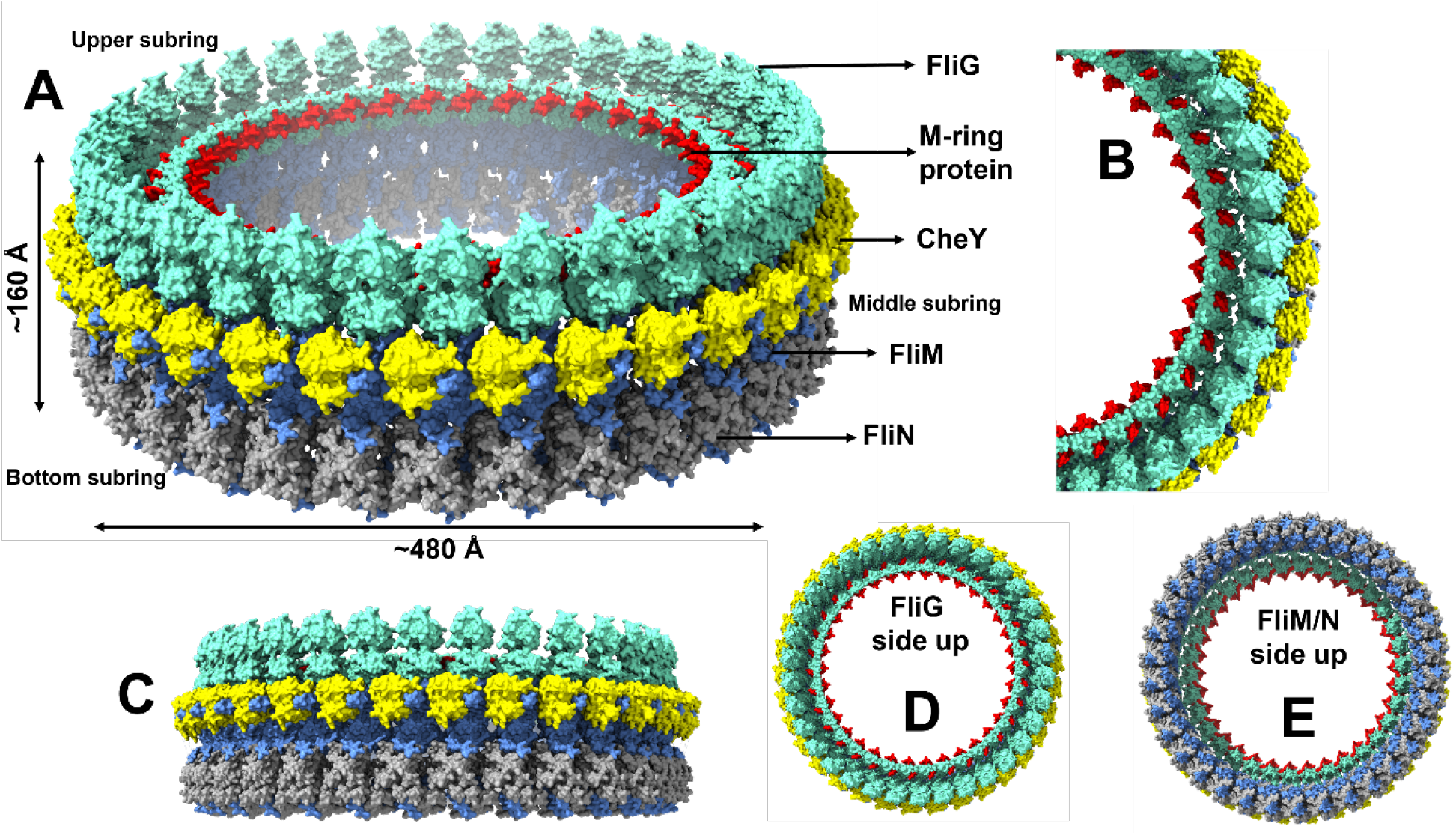
Cyro-EM ring structure of the flagellar C-ring of Salmonella enterica subsp. enterica serovar Typhimurium str. **LT2** (PDB ID: 8WIW): A. The large labelled structure on the top shows ~25 to 34 FliG monomers associating with roughly 34 M-ring protein (FliF_c_) monomers. FliG is in contact with CheY (in the CW mode), which in turn interacts with ~ 34 FliM and ~102 FliN. B. An inset showing a partial structure of the C-ring with colours retained as in panel A. Panel C – a side-view and D & E panels show the view from the top (upper subring) and bottom subring, respectively.

**Figure 5.**
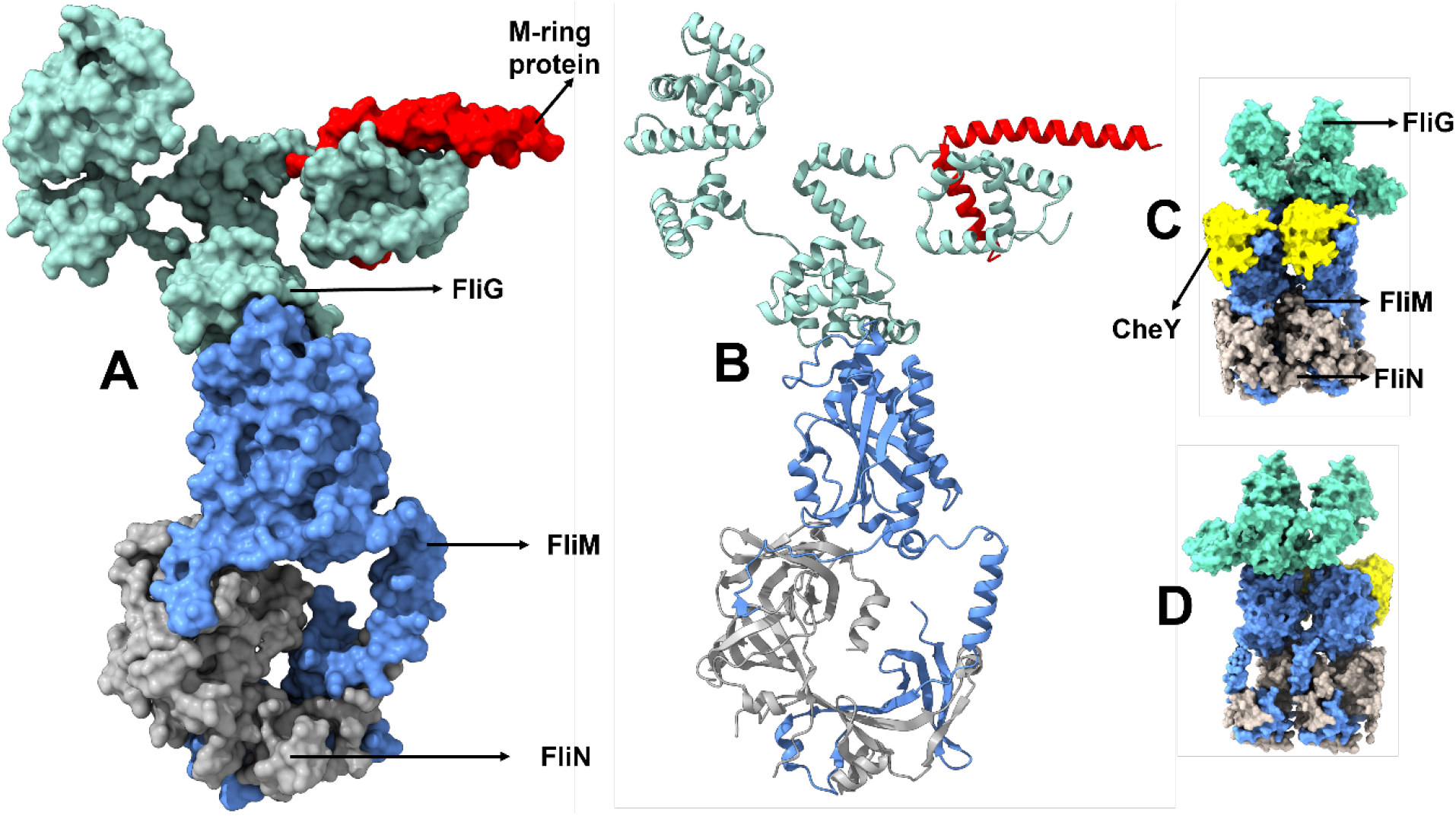
C-ring proteins of Salmonella enterica Typhimurium flagellar C-ring. (PDB ID: 8UMD). **A**. The basic unit of the flagellar C-ring comprising of interactions between M-ring protein (FliF_c_), FliG, FliM and FliN. Panel **B** shows the ribbon structures of the basic unit. Panel **C** and **D** show how two different fundamental units of the C-ring associate with each other.

### II.5 The reality of type 3 secretion system’s homology with BFS

The flagellar basal body is structurally and genetically homologous to the type 3 secretion system (T3SS) in bacteria and type 4 secretion system in archaea, both being molecular syringes that inject proteins/cellular contents into host cells or other bacteria. A comparison of the membrane-based protein assemblies associated with motility and secretion (with radial symmetry) found across the microbes can be seen in Table 3.

**Table 3.**
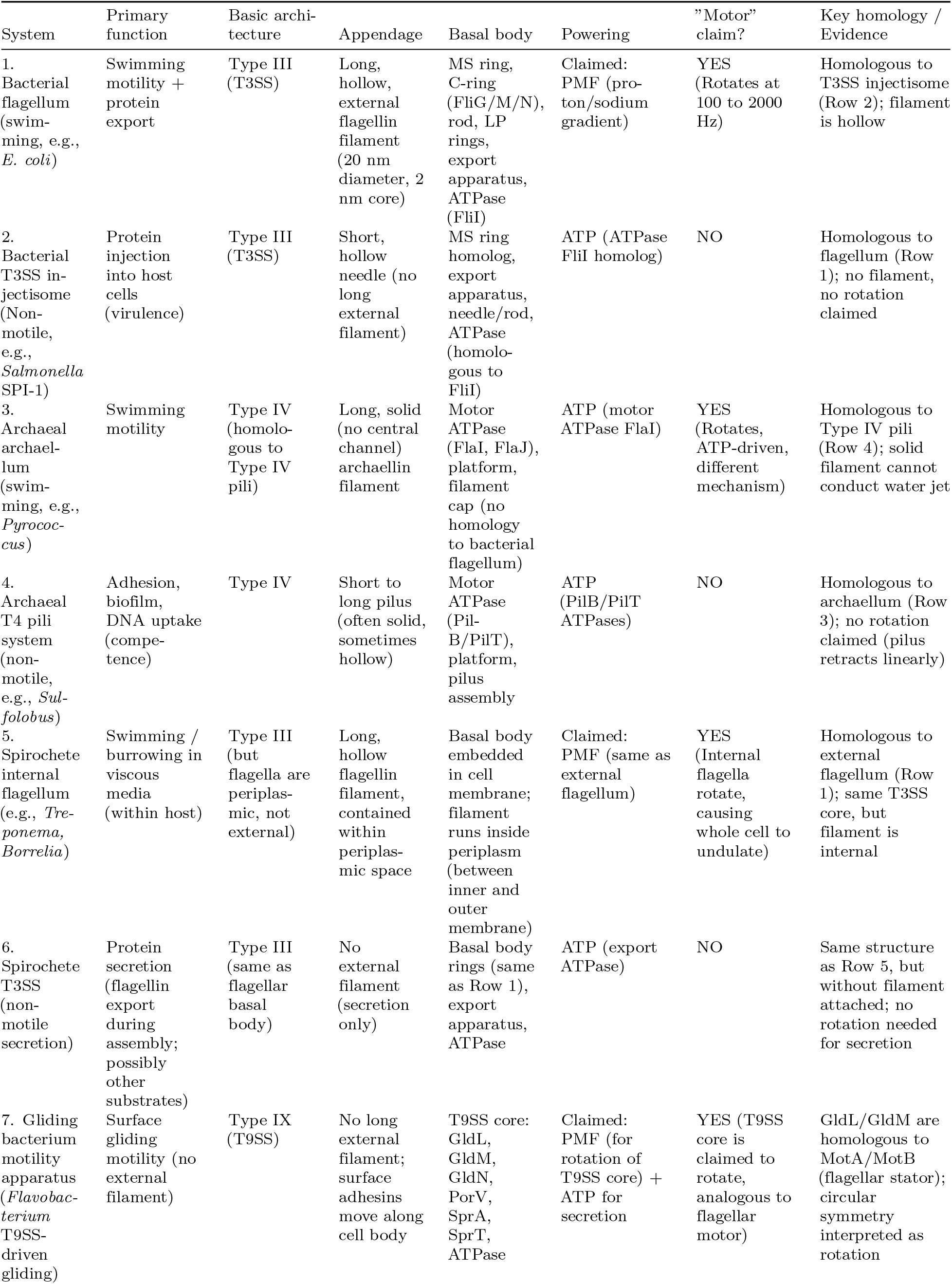
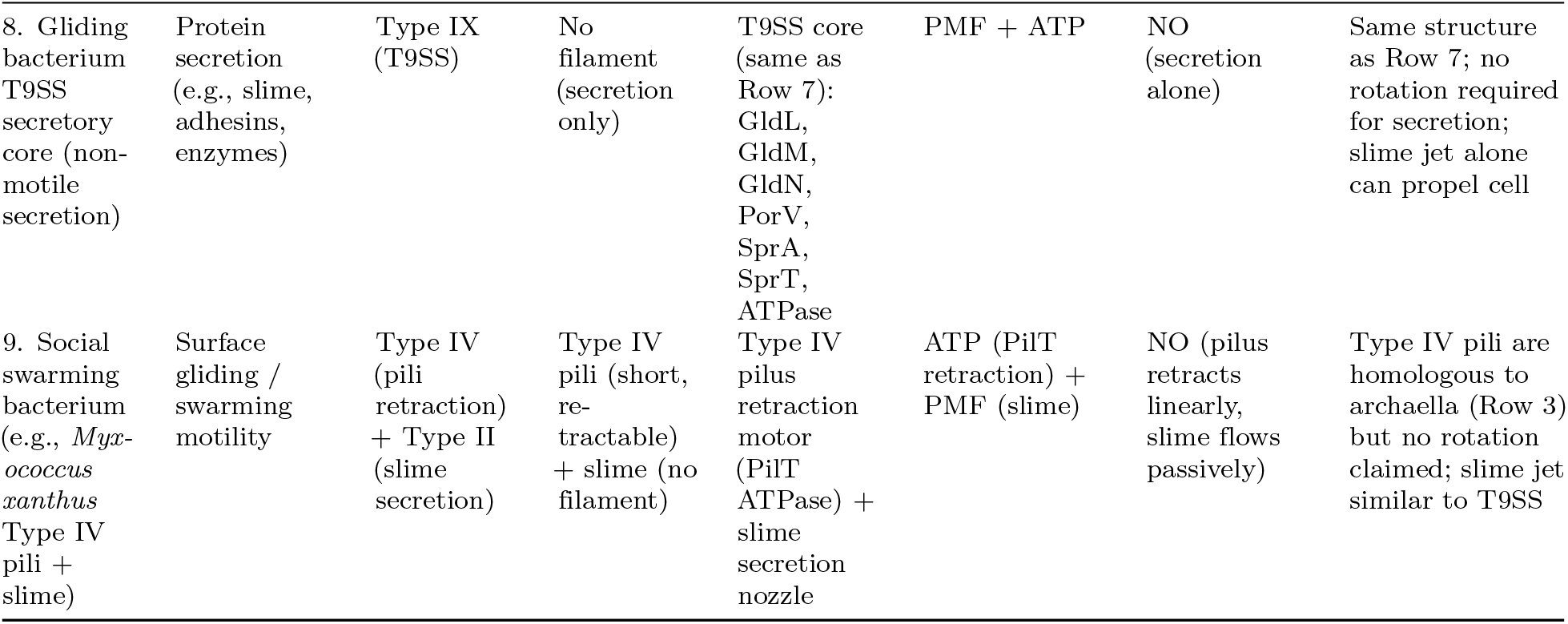
Comparison of motility and secretory apparatuses across four microbial systems.

| System | Primary function | Basic architecture | Appendage | Basal body | Powering | "Motor" claim? | Key homology / Evidence |
| --- | --- | --- | --- | --- | --- | --- | --- |
| 1. Bacterial flagellum (swimming, e.g., *E. coli*) | Swimming motility + protein export | Type III (T3SS) | Long, hollow, external flagellin filament (20 nm diameter, 2 nm core) | MS ring, C-ring (FliG/M/N), rod, LP rings, export apparatus, ATPase (FliI) | Claimed: PMF (proton/sodium gradient) | YES (Rotates at 100 to 2000 Hz) | Homologous to T3SS injectisome (Row 2); filament is hollow |
| 2. Bacterial T3SS injectisome (Non-motile, e.g., *Salmonella* SPI-1) | Protein injection into host cells (virulence) | Type III (T3SS) | Short, hollow needle (no long external filament) | MS ring homolog, export apparatus, needle/rod, ATPase (homologous to FliI) | ATP (ATPase FliI homolog) | NO | Homologous to flagellum (Row 1); no filament, no rotation claimed |
| 3. Archaeal archaelum (swimming, e.g., *Pyrococcus*) | Swimming motility | Type IV (homologous to Type IV pili) | Long, solid (no central channel) archaellin filament | Motor ATPase (FlaI, FlaJ), platform, filament cap (no homology to bacterial flagellum) | ATP (motor ATPase FlaI) | YES (Rotates, ATP-driven, different mechanism) | Homologous to Type IV pili (Row 4); solid filament cannot conduct water jet |
| 4. Archaeal T4 pili system (non-motile, e.g., *Sulfolobus*) | Adhesion, biofilm, DNA uptake (competence) | Type IV | Short to long pilus (often solid, sometimes hollow) | Motor ATPase (PilB/PilT), platform, pilus assembly | ATP (PilB/PilT ATPases) | NO | Homologous to archaelum (Row 3); no rotation claimed (pilus retracts linearly) |
| 5. Spirochete internal flagellum (e.g., *Treponema*, *Borrelia*) | Swimming / burrowing in viscous media (within host) | Type III (but flagella are periplasmic, not external) | Long, hollow flagellin filament, contained within periplasmic space | Basal body embedded in cell membrane; filament runs inside periplasm (between inner and outer membrane) | Claimed: PMF (same as external flagellum) | YES (Internal flagella rotate, causing whole cell to undulate) | Homologous to external flagellum (Row 1); same T3SS core, but filament is internal |
| 6. Spirochete T3SS (non-motile secretion) | Protein secretion (flagellin export during assembly; possibly other substrates) | Type III (same as flagellar basal body) | No external filament (secretion only) | Basal body rings (same as Row 1), export apparatus, ATPase | ATP (export ATPase) | NO | Same structure as Row 5, but without filament attached; no rotation needed for secretion |
| 7. Gliding bacterium motility apparatus (*Flavobacterium* T9SS-driven gliding) | Surface gliding motility (no external filament) | Type IX (T9SS) | No long external filament; surface adhesins move along cell body | T9SS core: GldL, GldM, GldN, PorV, SprA, SprT, ATPase | Claimed: PMF (for rotation of T9SS core) + ATP for secretion | YES (T9SS core is claimed to rotate, analogous to flagellar motor) | GldL/GldM are homologous to MotA/MotB (flagellar stator); circular symmetry interpreted as rotation |
| 8. Gliding bacterium T9SS secretory core (non-motile secretion) | Protein secretion (e.g., slime, adhesins, enzymes) | Type IX (T9SS) | No filament (secretion only) | T9SS core (same as Row 7): GldL, GldM, GldN, PorV, SprA, SprT, ATPase | PMF + ATP | NO (secretion alone) | Same structure as Row 7; no rotation required for secretion; slime jet alone can propel cell |
| 9. Social swarming bacterium (e.g., *Myxococcus xanthus* Type IV pili + slime) | Surface gliding / swarming motility | Type IV (pili retraction) + Type II (slime secretion) | Type IV pili (short, re-tractable) + slime (no filament) | Type IV pilus retraction motor (PilT ATPase) + slime secretion nozzle | ATP (PilT retraction) + PMF (slime) | NO (pilus retracts linearly, slime flows passively) | Type IV pili are homologous to archaella (Row 3) but no rotation claimed; slime jet similar to T9SS |

The classical model arbitrarily labels certain systems as “rotary motors” (rows 1, 3, 5, 7) based on circular symmetry, external appendages, and inferred analogies. Yet structurally homologous systems with identical core components (rows 2, 4, 6, 8, 9) are never claimed to rotate. The only consistent difference is the presence of a long filament (hollow or solid) or a slime nozzle. The classical model requires four independent mechanisms for the various microbes above: rotary flagellum (bacteria, H^+^/Na^+^-driven), archaellum (ATP-driven, different filament composition, not hollow), spirochete internal flagella (different geometry), and gliding machinery (different proteins, no flagella). As earlier (in Gideon et al., 2024, Section 4), we argue with concrete comparative evidence in Table 3 that the mere presence of a radial symmetry is not conclusive or indicative of rotary functionalism. A parsimonious explanation for motility across systems is that none of their basal modules need to rotate and all these are mere secretion systems. These radially arranged proteins on the membranes evolved to inject/secrete/eject water and other cellular components, and the filament/pili evolved to guide the ejected fluid for thrust (quite like the long tail of the kite stabilizes its flight, but only more roles in the low Re regime for bacterial motion). This evolutionary parsimonious take avoids the direct attack of irreducible complexity. A systematic study of Table 3 should convince the unbiased reader that: (i) every motility system is built on a secretory scaffold, (ii) secretory scaffolds can and need to function without rotation (T3SS, Type IV pili, T9SS core), (iii) the only difference between “rotary” and “non-rotary” systems is the presence of a long filament or slime nozzle and the perception in the eyes of the beholders (which was misdirected by the optical illusion of rotation!), (iv) rotation claims are inconsistent with structural features, applied only when circular symmetry is misinterpreted as a gear mechanism.

### II.6 The Zhou et al. (1998) mutation study and its implications

Before the 21^st^ century, the scientific consensus, built on structural and genetic evidence, pointed strongly to the C-ring (composed of FliG, FliM, and FliN proteins) as the rotor. The leading model of the time described a mechanism that relied on the rotational symmetry of the C-ring (Thomas et al., 1999). The stator complex (MotA and MotB) was seen as a stationary, ion-conducting channel that would interact with the rotating C-ring to generate torque. This model was so compelling that the discovery of any essential, conserved, protonatable residue for torque generation was expected to be found within the rotor proteins (FliG, FliM, FliN). Zhou et al. (1998) systematically tested this hypothesis by mutating every conserved acidic residue in the five proteins then thought to be involved in torque generation (MotA, MotB, FliG, FliM, FliN). The results were clear and surprising:

The negative result: Mutations to the conserved acidic residues in MotA, FliG, FliM, and FliN did not impair torque generation. This directly contradicted the expectation that the rotor contained the essential proton-binding sites.

The positive result: Only one residue proved essential for torque generation: aspartate 32 (D32) in the stator protein MotB. The paper concluded that “Asp 32 of MotB functions as a proton-binding site in the bacterial flagellar motor and that no other conserved, protonatable residues function in this capacity.” But as shown in Gideon et al. (2024) (Table 4), this residue is ALSO not conserved across motile flagellated bacteria and we could conclude that even that residue cannot be aiding any purported rotary function.

**Table 4.** Approximate metabolic oxidation stoichiometry of select six-carbon biological fuels.

| Substrate | Net equation |
| --- | --- |
| Glucose | $\text{C}_6\text{H}_{12}\text{O}_6 + 34 \text{ADPOH} + 34 \text{POH} + 6 \text{O}_2 \rightarrow 6 \text{CO}_2 + 34 \text{ATP} + 36 \text{H}_2\text{O}$ |
| Hexanoic acid | $\text{C}_6\text{H}_{12}\text{O}_2 + 46 \text{ADPOH} + 46 \text{POH} + 8 \text{O}_2 \rightarrow 6 \text{CO}_2 + 46 \text{ATP} + 56 \text{H}_2\text{O}$ |
| Leucine | $\text{C}_6\text{H}_{13}\text{NO}_2 + 24 \text{ADPOH} + 24 \text{POH} + 4 \text{O}_2 \rightarrow 6 \text{CO}_2 + \text{NH}_3 + 24 \text{ATP} + 24 \text{H}_2\text{O}$ |

Furthermore, the classical model has undergone a “chameleonic change” (Gideon et al., 2024, Section 1): Mot proteins were originally stators; now they are claimed to be the primary rotors (Santiveri et al., 2020; Wadhwa and Berg, 2022). This shift, occurring two decades after Zhou et al. s work, indicates the field lacks a stable mechanistic model. The classical view relies heavily on visual analogies and simplistic biophysical models, presents fundamental flaws in thermodynamics, structural mechanics, and energy efficiency. The realities of bacterial flagella-assisted motility field, when summated is:

1. direct visualization of nanoscale axial rotation is lacking,
2. physiological rotational frequencies are fully inferred,
3. passive helical dynamics can mimic rotational behavior,
4. interpretation is model-dependent and bacteria without flagella and with truncated flagella also show mobility.
5. circular structures are classical secretory system signs, and the flagellar system also has secretory function.
6. pmf-ionic differential based driving forces and transduction mechanisms are purely inferential, falling apart at the slightest inquiry
7. F_0_F_1_-ATPase was deemed a rotary motor partly because *γ*-subunit rotation was visualized directly with fluorescent probes (Noji et al., 1997) in a highly modified experimental setup. No equivalent direct visualization of the flagellar C-ring or rod rotation exists at atomic resolution in situ during function. (Also, the Complex V’s ATPase and ATPsynthase roles have been described in thermodynamically viable and non-reversible ways with the murburn model, questioning the veracity/applicability of the Noji experiment for Complex V.)

What is needed to explain the functionality of the structure is a stochastic operational mechanism, one which does not need any specific arrangement or any exact residue located at a particular locus of the protein to be conserved (quite unlike the deterministic rotational model) but only requires a preponderance of charged/polar residues at the proteins that line the interface) explains why the flagella system wasn ‘t’ selected’ out by natural selection over the course of evolution.

## III. The murburn model of bacterial motility: a parsimonious and quantitative framework

### III.1 Basics of formulation

We derive the governing equations from first principles of low-Reynolds-number fluid dynamics, slender-body theory, and active filament mechanics. The framework demonstrates that a metabolically sustained water ejection rate of *Q* ≈ 2× 10^−20^ m^3^*/*s through a sub-nanometre nozzle generates sufficient local shear 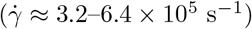 to excite a passive bending wave on a flexible, spirally grooved filament. The resulting traveling-wave propulsion, governed by the sperm number (Sp~ 1–10), yields a swimming speed of ≈30–56 *µ*m*/*s with mechanical power ≈0.15 pW, all without requiring rigid axial rotation, nanoscale gearing, or ion-gradient-driven torque.

The core foundational postulates are:

1. **The basal assembly is an active secretion-coupled flow generator**. It functions primarily as a transmembrane redox and secretion module, not as a torque-generating rotor. Water is produced metabolically (via DROS chemistry) and ejected at the hook–rod junction. The ejected water flows as an *external sheath along the filament surface*; it need not pass through the filament lumen, which makes the mechanism equally applicable to solid archaella that lack a hollow core.
2. **The external filament is a flexible elastohydrodynamic coupler**. It does not rotate rigidly. Instead, it propagates bending waves and precesses in response to the local shear flow created by the ejected water, channelled by the spiral grooves on its surface.
3. **Propulsion emerges from anisotropic viscous drag**. At low Reynolds number (Re ~ 10^−4^), thrust arises from the difference between perpendicular and parallel drag coefficients of a waving filament, not from inertial pushing or screw mechanics.

These postulates reframe the problem from engineering mechanics (gears, shafts, rotors) to soft-matter physics (active filaments, Stokes flow, elastohydrodynamics).

### III.2 Physical regime: Stokesian dynamics

For a bacterium of length *L*_*b*_ ~ 2 *µ*m swimming at speed *U* ~ 10^−4^ m*/*s in water (*ρ* = 10^3^ kg*/*m^3^, *η* = 10^−3^ Pa · s):

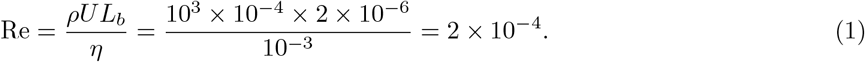

Thus Re ≪ 1: inertia is negligible and the Navier–Stokes equation reduces to the Stokes equation,

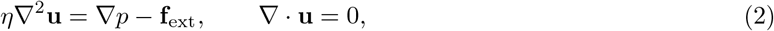

where **u** is fluid velocity, *p* pressure, and **f**_ext_ external forces. Consequences: forces balance instantaneously (no momentum conservation); reciprocal motion yields no net displacement (Scallop theorem); propulsion requires non-reciprocal deformation, satisfied by traveling waves or precession. Macroscopic screw and propeller analogies are invalid in this regime.

### III.3 Basal assembly as an active Stokesian source

The basal assembly (a Type III secretion system homolog) ejects water at a volumetric rate *Q*. Because the flow is ejected at the filament base, it creates a local shear field. The characteristic shear rate at the filament surface (radius *r*_*f*_ ≈ 10^−8^ m) is

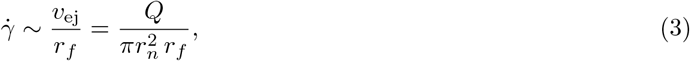

where 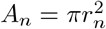 is the nozzle area (*r*_*n*_ ≈1 nm). For the metabolically sustained *Q* ≈ 2 ×10^−20^ m^3^*/*s (glucose oxidation, Section III.8):

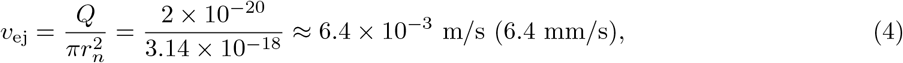

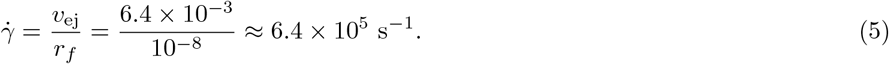

Such shear rates are more than sufficient to excite bending in a flexible filament (critical shear for flagellar buckling ~10^3^–10^4^ s^−1^). The basal assembly does not need to generate a high-speed jet; it only needs to produce a localized shear flow, and the filament does the rest. Structurally, the lateral exit of this water is plausible: the *Vibrio* hook and proximal filament present radial canal-like pores (~10 Å in diameter) surrounding the central channel (Fig. 7; Section IV), providing a candidate conduit for water to emerge along the basal region rather than solely from the distal tip.

#### Operating regime for *Q*

The quoted ranges already span the conservative-to-peak window rather than being pinned to the most favourable input: the steady-state glucose-oxidation estimate is *Q* ≈ 1 × 10^−20^ m^3^*/*s and the near-peak value ≈ 2 × 10^−20^ m^3^*/*s (Section III.8). Because 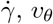 and *f*_precess_ all scale linearly with *Q*, this maps directly onto the precession band *f*_precess_ ≈1,800–3,600 Hz (lower end conservative, upper end near-peak). The propulsive amplitude *A* is geometric and *Q*-independent (Section III.6), so the predicted *U* ≈ 30–56 *µ*m*/*s is essentially insensitive to the choice of *Q* within this window. The shear that actually drives precession is reduced by the groove-transmission efficiency *η*_*t*_, which is carried explicitly through the equations of Section III.5; the observed precession in turn bounds it at *η*_*t*_ ≳ 0.5, so *η*_*t*_ enters the chain as a single, data-constrained factor rather than a free knob.

### III.4 Flagellum as an elastohydrodynamic filament

The flagellum is modeled as a thin, elastic, inextensible filament of length *L*_*f*_, radius *r*_*f*_, and bending rigidity *EI*, immersed in a viscous fluid. Its dynamics obey the forced elastohydrodynamic beam equation,

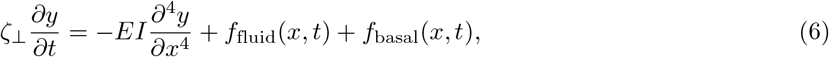

where *y*(*x, t*) is the transverse displacement, *ζ*_⊥_ ≈ 4*πη/* ln(*L*_*f*_ */r*_*f*_) the transverse drag coefficient per unit length, *f*_fluid_ the hydrodynamic forcing from the ambient flow, and *f*_basal_ the forcing from basal ejection (localized near *x* = 0). With *EI* ~ 10^−24^ N · m^2^ (from a bending persistence length of 5–10 *µ*m), *L*_*f*_ = 10^−5^ m, and *ζ*_⊥_ ≈1.82× 10^−3^ Pa· s, the natural bending dynamics are solved in Section III.5; the filament oscillates in the tens-of-Hz range when perturbed, with no external high-speed rotation required.

### III.5 Shear-induced transverse forcing, precession, and wave propagation

A central question is how an *axial* sheath flow excites a *transverse* bending wave. The resolution is geometric: the filament is helical, carrying spiral grooves from the supercoiled flagellin polymer. Water ejected at the hook–rod junction flows along the filament axis 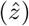, but because the local surface is inclined at the helix pitch angle *ψ*, the grooves convert axial flow into a circumferential component,

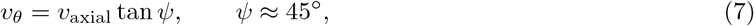

which produces a transverse drag force per unit length,

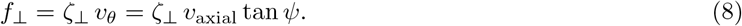

This bends the filament and drives precession of the bending plane. The hook bend enhances, but is not required for, this effect, consistent with hookless archaella remaining motile. The geometric coupling coefficient is

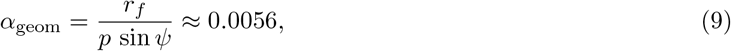

with helix pitch *p* ≈ 2.5 *µ*m, replacing a borrowed empirical estimate *α* ~ 0.01–0.1 and lying within a factor of two of it. The precession frequency follows, carrying the groove-transmission efficiency *η*_*t*_ (defined below) explicitly through the chain,

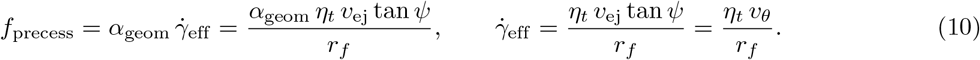

For order-unity transmission (*η*_*t*_ → 1) over the operating range *Q* ≈ 1–2 × 10^−20^ m^3^*/*s this gives 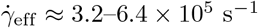 and hence *f*_precess_ ≈ 1,800–3,600 Hz; written this way, the observed precession directly *bounds η*_*t*_ (next paragraph) rather than leaving it unconstrained. This is the rate at which the bending plane rotates; it is distinct from the much slower frequency at which the beam itself flexes (Section III.8).

#### Axial-flow persistence

The expression for 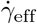 above assumes the axial flow stays close to *v*_ej_ over the forcing region. A *free* submerged stream could not achieve this: at Re~ 10^−5^ the ejected water carries negligible inertia, so as an unconfined Stokes source its centreline speed would fall as *v* ~*Q/*(4*πz*^2^) (a drop of ~10^9^ over *L*_*f*_), and persistence cannot come from “jetting.” It comes instead from *confinement* : the no-slip filament surface and the helical grooves channel the ejected water as a wall-bound creeping film, for which mass conservation (not inertia) sets the axial speed, *v*_axial_ = *η*_*t*_ *Q/a*_*g*_ ≈ *η*_*t*_ *v*_ej_ (taking the groove cross-section *a*_*g*_ comparable to the nozzle), with the cumulative leakage along the channel absorbed into a transmission efficiency *η*_*t*_ ≤ 1. The effective shear is then 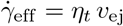 tan *ψ/r*_*f*_, so that *f*_precess_ ∝ *η*_*t*_.

Motility data bound *η*_*t*_ tightly, and from the favourable direction: the measured “rotation” of the *Vibrio* polar flagellum reaches ≈1.7 kHz (Magariyama et al., 1994), which requires *η*_*t*_ ~*≪*𝒪 (1) (≳ 0.5), whereas free-jet dispersal (*η*_*t*_ ~10^−9^) would imply a precession many orders of magnitude too slow to register. The observed kHz precession therefore *excludes* the free-jet objection and is consistent with groove-confined flow transmitted at order-unity efficiency over the filament length. We are careful not to overstate the mechanism here: the confinement picture (a wall-bound creeping film guided by the helical grooves) is a physical argument that the motility data render plausible and internally self-consistent, but that has not yet been imaged directly. A resolved measurement (or numerical-hydrodynamics treatment) of the near-surface film is the natural experimental test and is left to future work; what the present data establish unambiguously is the *bound η*_*t*_ ≳ 0.5, which is all the quantitative chain requires.

#### Groove coupling and the no-slip surface

It is natural to worry that no-slip and leakage at the groove walls might degrade the axial→ azimuthal conversion. In fact the no-slip condition is the *origin* of the coupling, not a loss. A smooth, perfectly slipping cylinder would let the ejected film slide axially without acquiring any azimuthal component (*v*_*θ*_ → 0); the no-slip, anisotropically grooved flagellin surface instead presents direction-dependent drag, the same anisotropy (*ξ*_⊥_ *> ξ*_∥_) that underlies resistive-force theory (Section III.6), so that an axial flow over the helically wound grooves necessarily develops the transverse component *v*_*θ*_ = *v*_axial_ tan *ψ*. This coupling vanishes on a slippery surface and strengthens with groove depth; for the deeply grooved flagellin lattice it is of order unity. The directional conversion therefore introduces *no* additional loss beyond the transmission factor *η*_*t*_ already carried in 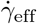; equivalently, an effective coupling *α*_eff_ = *η*_*t*_ *α*_geom_ ≈ (0.5–1) × 0.0056, so a single, data-constrained efficiency (*η*_*t*_ ≳ 0.5, fixed above by the observed rotation rate) governs the entire shear-to-precession chain. A first-principles value of *η*_*t*_ from a detailed numerical-hydrodynamics treatment of flow over the flagellin surface would sharpen this estimate but is not required for the present results.

### III.6 Propulsion via anisotropic drag (resistive-force theory)

Once the filament propagates a traveling wave of amplitude *A* and wavenumber *k* = 2*π/λ*, it generates thrust through anisotropic viscous drag. For a slender filament the drag per unit length is **F**_drag_ = −*ξ*_∥_**u**_∥_ −*ξ*_⊥_**u**_⊥_ with *ξ*_∥_ ≈2*πη/* ln(*L*_*f*_ */r*_*f*_) and *ξ*_⊥_ ≈2*ξ*_∥_. For a traveling wave *y*(*x, t*) = *A* sin(*kx* −≪*ωt*), resistive-force theory (Gray and Hancock, 1955; Lighthill, 1976) gives the net swimming speed

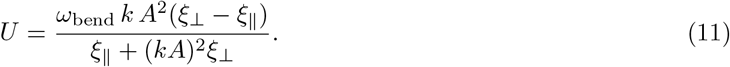

Using the wave geometry derived below (*A* ≈ 0.25 *µ*m, consistent with the geometric value *p* sin *ψ/*2*π* ≈ 0.28 *µ*m; *n*_waves_ ≈ 1.5, so *λ* = *L*_*f*_ */n*_waves_ ≈ 6.7 *µ*m, *k* ≈ 9.4 × 10^5^ m^−1^), *ω*_bend_ ≈ 4.9 × 10^2^ rad*/*s, and *ξ*_⊥_*/ξ*_∥_ ≈ 2, we obtain a thrust *F*_thrust_ ≈ 0.8 pN and

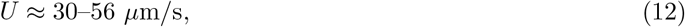

within the observed swimming range. The two ends of this band are the two natural resistive-force estimates: the lower end (≈ 30 *µ*m*/*s) is the force-free self-propulsion speed given directly by the expression above, while the upper end (≈56 *µ*m*/*s) follows from balancing the thrust *F*_thrust_ ≈0.8 pN against the cell-body drag (*ζ*_body_ ≈1.4 × 10^−8^ N s*/*m, so *U* = *F*_thrust_*/ζ*_body_). They differ by the usual resistive-force prefactor and amplitude uncertainty, and both lie within the measured range. Single-polar-flagellum *Vibrio alginolyticus* cells swim at tens of *µ*m*/*s, rising to *>*100 *µ*m*/*s for the fastest cells; Magariyama et al. (1995) measured a roughly *linear* speed–rotation relation (ratio *U/f* ≈0.113 *µ*m) that *saturates* at high rotation rate (Atsumi et al., 1996). The predicted *U* ≈30–56 *µ*m*/*s sits squarely within this measured band, and the observed saturation is exactly the behaviour implied by the geometric, *Q*-independent wave amplitude derived below (Section III.6), and is inconsistent with an unbounded *U*∝ *Q*^2^ scaling. This expression does not depend on the filament rotating rigidly; it requires only a propagating bending wave.

#### Wave geometry from the beam balance

The amplitude and wave count need not be borrowed from observation; both are constrained by the forced beam dynamics. *Amplitude*. In steady precession the filament’s off-axis excursion *A* sweeps azimuthally at 2*πf*_precess_, so its transverse material velocity *A* (2*πf*_precess_) is set by the driving groove-flow velocity *v*_*θ*_ = *η*_*t*_*v*_ej_ tan *ψ*. Equating the two and inserting *f*_precess_ = *α*_geom_*η*_*t*_*v*_ej_ tan *ψ/r*_*f*_ (Section III.5), the efficiency *η*_*t*_ and the ejection rate *cancel* explicitly, leaving a purely geometric amplitude,

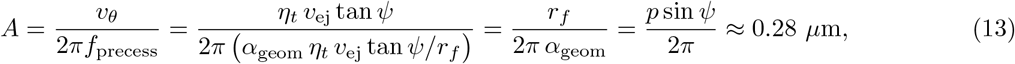

essentially the supercoil radius, in good agreement with the observed *A* ≈ 0.25 *µ*m and, importantly, independent of the uncertain *η*_*t*_ and *Q. Wave count*. The sperm number obeys Sp = *L*_*f*_ (*ζ*_⊥_*ω*_bend_*/EI*)^1*/*4^ = *kL*_*f*_ = 2*πn*_waves_, so the wave count is not an independent parameter: *n*_waves_ = Sp*/*2*π*. The elastohydrodynamic optimum for propulsion, Sp ≈ 4–10, therefore fixes *n*_waves_ ≈ 0.6–1.6, bracketing the value 1.5 used above. Thus *A* is a geometric *prediction* and *n*_waves_ is pinned to order unity by the efficiency optimum rather than fitted; *U* and *F*_thrust_ inherit this status. A full boundary-value solution of the forced Euler–Bernoulli equation would fix the *O*(1) prefactor of *n*_waves_ exactly and is left to future work.

### III.7 The sperm number and efficient propulsion

The dimensionless sperm number compares viscous to elastic forces:

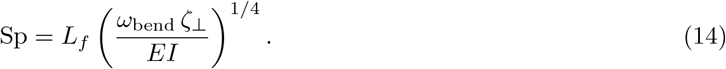

With *L*_*f*_ = 10^−5^ m, *ω*_bend_ ≈ 4.9 × 10^2^ rad*/*s (the bending-relaxation rate that sets the power and thrust, *not* the kHz precession frequency), *ζ*_⊥_ ≈ 1.82 × 10^−3^ Pa · s, and *EI* ≈ 10^−24^ N · m^2^, one obtains Sp ≈ 9.7, at the upper end of the optimal ~1–10 range for efficient propulsion. Equivalently Sp = *kL*_*f*_ = 2*πn*_waves_ (Section III.6), so the filament is mechanically tuned to convert basal shear into propulsive waves.

### III.8 Water budget, frequency decoupling, and energetics

#### Water production

Metabolism is a copious source of water. A minimalistic combustion stoichiometry of the basic six-carbon building blocks of carbohydrates, lipids and proteins (glucose, hexanoic acid and leucine) gives

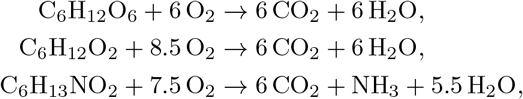

i.e. roughly one equivalent each of carbon and molecular oxygen yields about one equivalent each of CO_2_ and water by direct combustion. Physiologically the route differs; taking glucose as the worked example, routine enzymatic metabolism gives

i. routine enzymatic metabolism gives

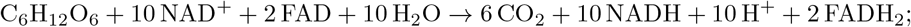
ii. murburn oxidative phosphorylation of NADH/FADH_2_ (approximate stoichiometry 3:2) gives

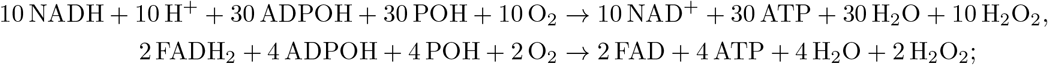

and (iii) catalase/oxidase activity converts the peroxide to water, 12 H_2_O_2_→ 12 H_2_O + 6 O_2_. The net mass- and charge-balanced equation for glucose is therefore

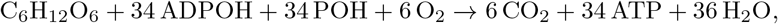

i.e. 36 water molecules per glucose; Table 4 lists the net equations for the other fuels.

At a metabolic rate of ~ 10^7^ glucose/s the water production is

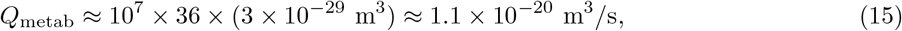

consistent with the *Q* ≈2 × 10^−20^ m^3^*/*s used above; the ejected water is thus sustained directly by metabolism.

#### Water homeostasis and the scallop theorem

The bacterial membrane carries aquaporins at a density of ~10^4^ *µ*m^−2^ that transport water bidirectionally (Verkman, 2013), so any transient lowering of cytosolic water is replenished through them during the intervals when the flagellar module is inactive. Because a cell swims continuously for only ~ 1 s (up to ~ 10 s under strongly favourable chemotaxis, when fuel is not limiting), water availability is not a limiting factor: the alternation between brief swimming bouts and metabolic / aquaporin-mediated replenishment does not conflict with the scallop theorem.

#### Two distinct rates, and why the budget is robust to which one dominates

The motion involves two physically distinct frequencies that must not be conflated. The *precession frequency f*_precess_ ≈ 1.8–3.6 kHz (Section III.5) is the rate at which the bending plane sweeps about the filament axis, the kinematic signature an observer reads as “rotation.” The *bending relaxation rate ω*_bend_ is the intrinsic elastohydrodynamic response rate of the overdamped filament. Being inertia-free, the filament obeys the hyperdiffusive relation 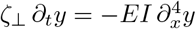, whose wavenumber-dependent relaxation rate (Wiggins and Goldstein, 1998) is

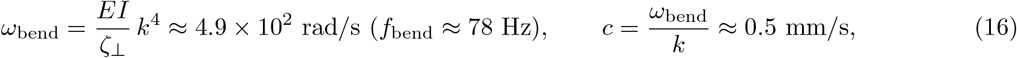

for *λ* = *L*_*f*_ */n*_waves_ ≈ 6.7 *µ*m (*k* ≈ 9.4 × 10^5^ m^−1^). The filament thus behaves as a mechanical low-pass filter: the large-amplitude, thrust-producing deformation is built at the slow relaxation rate *ω*_bend_, while the fast precession merely re-orients an already-formed bend.

#### Mechanical power

The viscous dissipation of the waving filament, 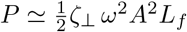, evaluated at these two rates brackets the mechanical cost:

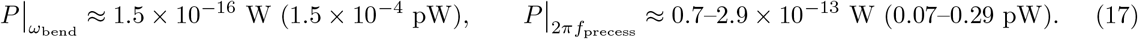

Taking a single representative (precession-frequency) value, *P* ≈ 0.15 pW: for a single polar flagellum (e.g. *Vibrio*) this is *<*1% of the cellular metabolic budget (~19–24 pW), and even at the highest precession frequency in the band (*P* ≈ 0.29 pW) the cost stays at ≈ 1.5%; four flagella (≲ 0.6–1.2 pW) remain at the few-percent level. Crucially, the budget holds across the *entire* range (whether the dissipation is charged to the fast precession or the slow flexing), so the energetic viability does *not* hinge on a delicate frequency assignment. A naively high precession frequency (≈ 12.7 kHz, i.e. *ω* ≈ 8 × 10^4^ rad*/*s) would inflate *P* by ~30× to≈ 4.6 pW; the geometrically derived *f*_precess_ = 1.8–3.6 kHz keeps the cost negligible. This range is, moreover, consistent with the fastest directly measured flagellar “rotation”, up to ≈1.7 kHz for the *Vibrio* polar flagellum (Magariyama et al., 1994), which the present model reinterprets as bending-plane precession rather than rigid axial rotation. It is worth being explicit about what such assays measure: tethered-cell and bead-rotation experiments register the periodic azimuthal signature of the appendage, which in this model is precisely the precession of the bending plane at *f*_precess_, not a rigid axial spin of the filament. The reinterpretation is therefore of the *mechanism* behind the observed rate, not of the rate itself, so no conflict with the measured frequencies arises.

### III.9 Force-balance comparison: jet propulsion, rigid rotation, and elastohydrodynamics

The three candidate mechanisms obey the *same* low-Reynolds-number force balance and differ only in how thrust is powered. Evaluated against the corrected chain (Table 5), only the elastohydrodynamic route is simultaneously sufficient and metabolically sustainable.

**Table 5.** Force-balance comparison of the three propulsion mechanisms. (corrected *Vibrio* chain). All obey the same low-Reynolds-number drag balance; they differ only in the power source.

| Mechanism | Thrust | Power / resource | Verdict |
| --- | --- | --- | --- |
| Direct jet | $\rho Q v_{\text{ej}} \approx 1.3 \times 10^{-19} \text{ N}$<br>($\sim 6 \times 10^6$ too small) | needs $Q \approx 5 \times 10^{-17} \text{ m}^3/\text{s}$;<br>empties cell in $\approx 0.02 \text{ s}$ | Excluded |
| Rigid helical rotation | $\approx 0.8 \text{ pN}$ (RFT) | rotary motor + PMF torque | Hydrodynamically OK; motor untenable (Secs. I–II) |
| Murburn elasto-hydrodynamic | $\approx 0.8 \text{ pN}$ (RFT) | shear from sustainable $Q \approx 2 \times 10^{-20} \text{ m}^3/\text{s}$; $< 1\%$ budget | Sufficient & sustainable |

#### Direct jet propulsion is excluded on two independent grounds

In the Stokes regime there is, to leading order, no inertial reaction thrust at all (Spagnolie and Lauga, 2010); even the generous inertial estimate *F*_jet_ = *ρQv*_ej_ ≈1.3 ×10^−19^ N falls short of the required ≈0.8 pN by a factor ~6× 10^6^. Worse, obtaining 0.8 pN by jetting would demand *Q* ≈ 5 ×10^−17^ m^3^*/*s, some 2500× the metabolic rate, draining the cell’s water content (~ 1 *µ*m^3^) in ≈0.02 s, physically impossible.

#### Rigid helical rotation is hydrodynamically adequate but mechanistically untenable

A rigid helix turning at kHz produces a comparable resistive-force thrust, but only if a rotary motor supplies the matching torque against viscous drag, the proton-motive stator–rotor machinery whose thermodynamic and structural implausibility is set out in Sections I–II.

#### The elastohydrodynamic mechanism succeeds with sustainable resources

The murburn route spends the modest, metabolically sustained *Q* ≈ 2 × 10^−20^ m^3^*/*s not on direct thrust but on generating *shear* (Section III.3), which the spiral grooving converts into a transverse bending wave; resistive-force theory then yields *F*_thrust_ ≈0.8 pN and *U* ≈ 30–56 *µ*m*/*s (Section III.6) at *<* 1% of the metabolic budget (Section III.8). The mechanism is, in effect, a shear-to-thrust *amplifier* : it reaches the required force with ~2500× less water than a jet and with no rotary motor at all. This is the quantitative backbone of the model: not that elastohydrodynamics is merely *a* viable option, but that, of the three, it is the *only* one meeting the thrust requirement with sustainable water and without a rotary engine.

### III.10 Summary of quantitative aspects

Table 6 collects the corrected, internally consistent values; Table 7 defines every symbol to keep the three distinct frequencies unambiguous.

**Table 6.** Murburn elastohydrodynamic model: corrected quantitative summary (*Vibrio*, single polar flagellum as worked case).

| Quantity | Symbol | Value | Source/Notes |
| --- | --- | --- | --- |
| Water ejection rate | $Q$ | $1\text{--}2 \times 10^{-20} \text{ m}^3/\text{s}$ | Glucose oxidation (III.8) |
| Nozzle radius | $r_n$ | 1 nm | Filament core |
| Ejection velocity | $v_{\text{ej}}$ | 3.2–6.4 mm/s | $Q/\pi r_n^2$ |
| Axial flow speed | $v_{\text{axial}}$ | $\approx \eta_t v_{\text{ej}}$ | $\eta_t \gtrsim 0.5$, confined groove flow (III.5) |
| Helix pitch angle | $\psi$ | $45^\circ$ | Literature |
| Circumferential speed | $v_\theta$ | $v_{\text{axial}} \tan \psi$ | Spiral grooves |
| Effective shear rate | $\dot{\gamma}_{\text{eff}}$ | $3.2\text{--}6.4 \times 10^5 \text{ s}^{-1}$ | $v_\theta/r_f$ |
| Geometric coupling | $\alpha_{\text{geom}}$ | $\approx 0.0056$ | $r_f/(p \sin \psi)$ |
| Precession frequency | $f_{\text{precess}}$ | 1,800–3,600 Hz | $\alpha_{\text{geom}} \dot{\gamma}_{\text{eff}}$ |
| Wavelength | $\lambda$ | 6.7 $\mu\text{m}$ | $L_f/n_{\text{waves}}$ |
| Bending frequency | $f_{\text{bend}}$ | $\approx 78 \text{ Hz}$ | Dispersion relation |
| Wave speed | $c$ | $\approx 0.5 \text{ mm/s}$ | $\omega_{\text{bend}}/k$ |
| Power per flagellum | $P$ | $\approx 0.15 \text{ pW}$ | $\frac{1}{2} \zeta_\perp \omega^2 A^2 L_f$ at $f_{\text{precess}}$; $< 1\%$ budget (III.8) |
| Thrust | $F_{\text{thrust}}$ | $\approx 0.8 \text{ pN}$ | Resistive-force theory |
| Swimming speed | $U$ | $\approx 30\text{--}56 \text{ }\mu\text{m/s}$ | RFT (self-propulsion to body-drag balance) |

**Table 7.** Symbol table. The three frequencies are physically distinct and must not be conflated.

| Symbol | Meaning |
| --- | --- |
| $Q$ | metabolic water ejection rate |
| $v_{\text{ej}}$ | ejection velocity at the nozzle ($Q/\pi r_n^2$) |
| $v_{\text{axial}}$ | axial sheath-flow speed along filament |
| $v_\theta$ | circumferential flow component from grooves |
| $\dot{\gamma}_{\text{eff}}$ | effective local shear rate at filament surface |
| $\alpha_{\text{geom}}$ | geometric shear-to-precession coupling ($r_f/p \sin \psi$) |
| $\eta_t$ | groove-flow transmission / coupling efficiency ($\gtrsim 0.5$) |
| $\alpha_{\text{eff}} = \eta_t \alpha_{\text{geom}}$ | effective (leakage-corrected) coupling |
| $f_{\text{precess}}$ | rate the bending plane rotates (kHz) |
| $f_{\text{bend}}, \omega_{\text{bend}}$ | rate the beam flexes (tens of Hz); sets dissipation |
| $A, \lambda, k$ | wave amplitude, wavelength, wavenumber |
| $L_f, r_f, r_n$ | filament length, filament radius, nozzle radius |
| $EI, \zeta_\perp, \xi_{\parallel, \perp}$ | bending rigidity, transverse drag, RFT drag coefficients |
| Sp | sperm number |

### III.11 Assumptions and limitations

The model above is quantitatively self-consistent. The four inputs that earlier stood as open assumptions have now each been addressed within the model (we recap them here, with the section where each is grounded); the only residual is an *O*(1) prefactor in the wave count, deferred to a future boundary-value solution.

#### Axial-flow persistence (addressed; Section III.5)

A free sub-nanometre jet would disperse as a Stokes source (*v* ~*Q/*4*πz*^2^, a ~10^9^ drop over *L*_*f*_) and could not persist; persistence instead follows from confinement of the ejected film by the no-slip surface and helical grooves, for which mass conservation sets the speed. The net transmission is carried by a single efficiency *η*_*t*_ ≤ 1, bounded ≳ 0.5 by the measured kHz “rotation” of the *Vibrio* flagellum, which by the same token *excludes* the free-jet objection (it would imply *η*_*t*_ ~ 10^−9^ and a precession orders of magnitude too slow to observe).

#### Groove coupling (addressed; Section III.5)

No-slip is the *origin* of the axial→azimuthal conversion, not a loss: the grooved surface’s drag anisotropy (*ξ*_⊥_ *> ξ*_∥_) generates *v*_*θ*_ = *v*_axial_ tan *ψ*, with effective coupling *α*_eff_ = *η*_*t*_ *α*_geom_ ≈ (0.5–1) × 0.0056. No additional loss arises beyond the single factor *η*_*t*_ above; a first-principles numerical value of *η*_*t*_ would refine, but not change, the picture.

#### Frequency budget (addressed; Section III.8)

The mechanical power is *<* 1% of the metabolic budget whether the dissipation is charged to the fast precession (≈ 0.15 pW) or the slow flexing (~ 10^−4^ pW). The result is therefore robust to the frequency assignment and does *not* hinge on a delicate decoupling; an inflated ≈ 4.6 pW figure would follow only from an unphysically high precession frequency (≈ 12.7 kHz).

#### Wave geometry (addressed; Section III.6)

The amplitude is now obtained from the beam balance, *A* = *p* sin *ψ/*2*π* ≈ 0.28 *µ*m (a geometric prediction, independent of *η*_*t*_ and *Q*, matching the observed ≈ 0.25 *µ*m), and the wave count is fixed to order unity by the propulsion optimum, *n*_waves_ = Sp*/*2*π* ≈ 0.6–1.6. *U* and *F*_thrust_ are therefore predictions up to the *O*(1) wave-count prefactor; a full boundary-value solution of the forced Euler–Bernoulli equation would fix that prefactor exactly and is the one remaining refinement.

#### Scope

We present *Vibrio* (single polar flagellum) as the rigorous worked case. Multi-flagella bundling (e.g. *E. coli*) and its hydrodynamic synergy are deferred to future work; the broad “microbial motility” claim is therefore demonstrated quantitatively for the unipolar case and argued qualitatively for the rest.

#### Illustrative animation

A short script that renders the closed-form relations above (the cell body translating at the predicted *U* while the filament carries the geometric wave) is provided in Supplementary

Appendix 3. It is a *kinematic visualisation for intuition only* : it draws the analytically derived motion and does *not* solve the fluid equations, is *not* a hydrodynamic simulation, and constitutes neither a validation nor a proof of the model. For visibility, the on-screen wave is deliberately slowed far below the physical kHz precession by a fixed display factor, so the animation’s apparent frequency is not the physical one.

## IV: Applications and testable predictions of the murburn model

The preceding sections have established the theoretical foundations of the murburn model for bacterial flagellar motility and detailed its mechanistic departures from the classical rotary paradigm. We now turn to the explanatory and predictive power of this framework. A scientific model gains credibility not merely by critiquing its predecessors but by accounting for diverse experimental observations, unifying disparate phenomena under a single principle, and generating falsifiable predictions that distinguish it from competing hypotheses.

### IV.1 Reinterpretation of key experiments & structural facets

#### Tethered cell experiments

When a filament is attached to a surface and the basal assembly ejects water, the cell body may counter-rotate slowly (~10 Hz) due to viscous torque from the asymmetric flow. This does not prove the filament rotates at kHz frequencies; it only shows that the basal ejection can create a net torque under non-physiological tethering.

#### Gold bead assays

Beads attached to filaments may trace circles or ellipses due to precession of the helical filament in the shear flow, not due to rigid axial rotation. The **Ali et al. (2016)** experiment clearly shows that a tethered filament (no basal motor) exhibits apparent rotation in a shear flow – exactly as predicted by the murburn model.

#### PMF-dependence?

This idea is irrelevant or redundant in the new theorization. Proton is used as a reactant, and not a cycling gradient or force. The fact that bacteria can swim well at alkaline pH quite well should be taken as conclusive evidence against the proton-cycling energetics.

##### 1. The hook as a “Universal Joint” – An overinterpretation

Das et al. (2026) **model the hook as a torque diverter that splits motor torque into revolution and spinning components. They introduce a Hill-type equation for hook stiffness (which is not worth reproducing here. The murburn critique views the hook differently:**

- The hook is not a mechanical universal joint; it is a flexible secretion guide for water and flagellin monomers.
- The “bending angle” *ψ* is not a torque-driven parameter but a passive hydrodynamic consequence of water ejection.
- The 33-fold stiffness increase from CCW to CW (*κ*_h_ from 100 to 3310 k_B_T) is an ad hoc fitting parameter, not based on any structural measurement.

##### 2. Statistical spread of hook morphology

Das et al. treat the hook as a fixed mechanical component with invariant properties. However, across bacterial species, the hook (FlgE polymer) shows enormous variability. Variations in hook length, hook diameter, FlgE monomer length as well as supercoiling state present logical questions against conservation of the hook in diverse bacteria (Table 8).

**Table 8.**
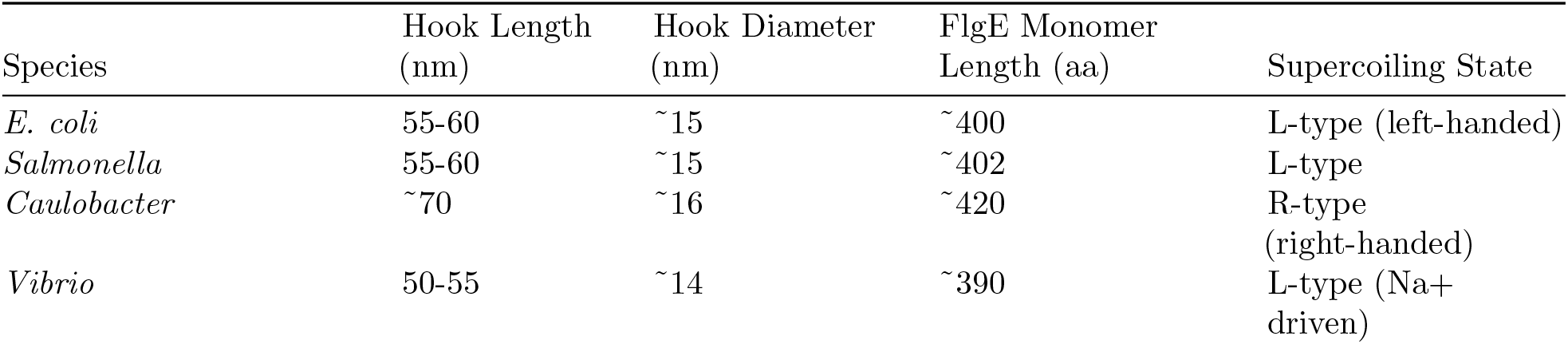
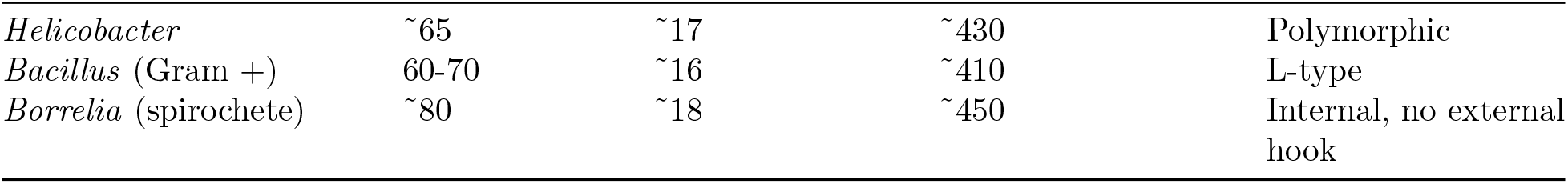
Comparison of the hook segment in diverse bacteria.

If the hook were a precision torque-transmitting universal joint, its dimensions and monomer composition would be highly conserved. They are not. The variation in length (50-80 nm), diameter (14-18 nm), and monomer length (390-450 aa) across species indicates that the hook s primary function is not mechanical torque transmission but flexible channeling of secreted material. From the cryo-EM structures of the bacterial flagellar axial proteins (Figures 6–8), these assemblies appear not as solid protein rods but as hydrated, porous structures whose internal water content and solvent connectivity may contribute to the hydrodynamic behaviour of the BFS. Re-examining publicly deposited cryo-EM structures of the flagellar filament (*Vibrio alginolyticus*, PDB 9RCB (Qin et al., 2026); *Helicobacter pylori*, PDB 9YGU (Kumar et al., 2026)) and of the flagellar hook (PDB 9M9F (Chen et al., 2025)) in ChimeraX, we find solvent-accessible inter-subunit pores of order 10 Å surrounding the central channel (Figure 7), recurring in a periodic, circumferential pattern along the subunit interfaces. Such inter-subunit gaps are consistent with prior structural reports that the flagellar axial assemblies retain ordered water in these spaces. We interpret them, alongside the well-characterised central channel, as candidate conduits for local water occupancy, exchange, and lateral streaming, providing a plausible structural pathway for the water transport invoked by the murburn mechanism. We stress that this is an interpretation of existing structures rather than a new structural determination.

**Figure 6.**
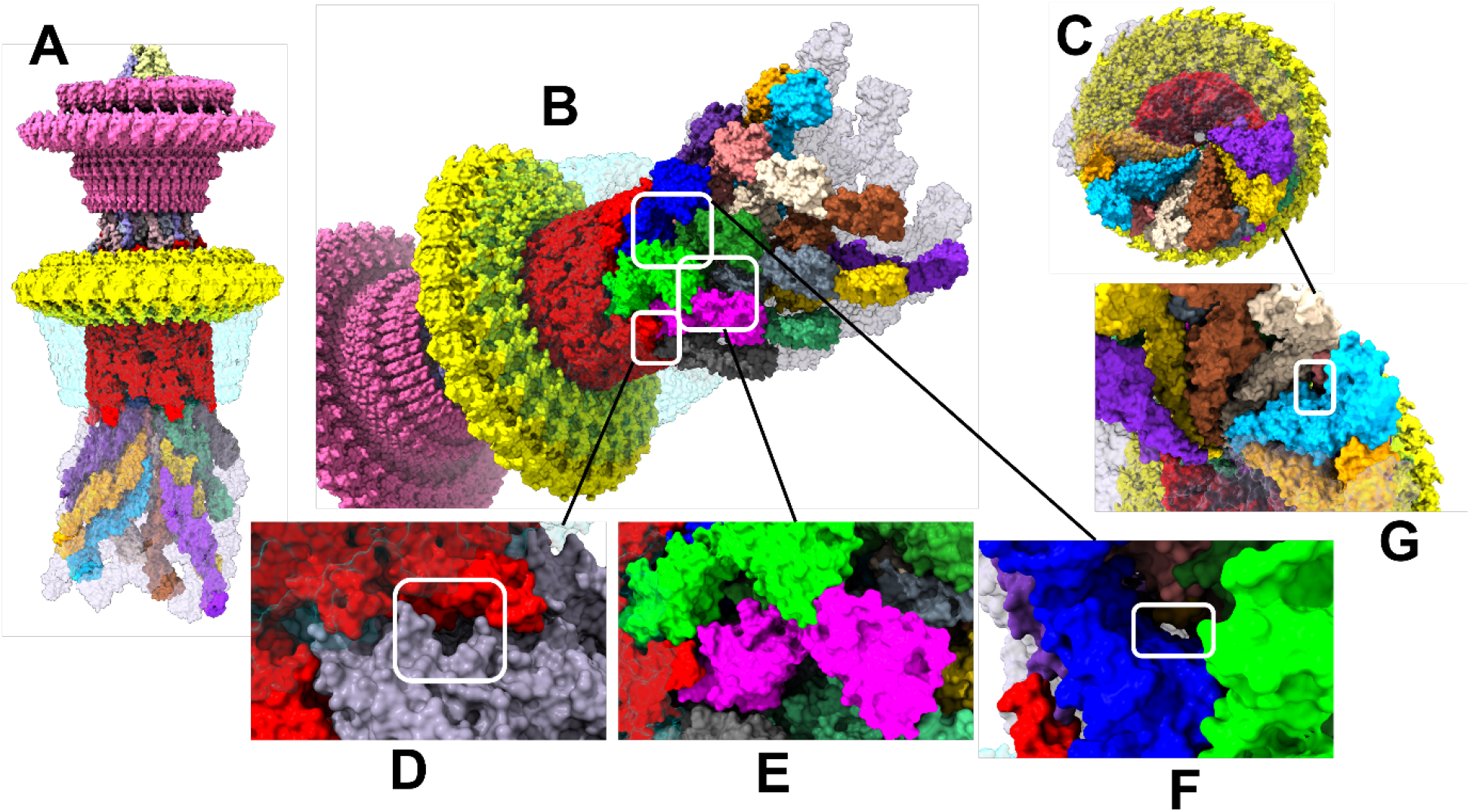
Bacterial flagellar basal body and hook. **A –** Structure of the Salmonella enterica subsp. enterica serovar Typhimurium str. LT2 hook (PBD ID: 7CGO) featuring LP ring, M-ring and flagellar protein assembly. For more details of the parts, refer to Figure 5 panel **A**. Nearly half of the flagellar protein monomers were coloured and the surface of the L-ring was made transparent to show the interaction of FlgF and FlgG with the flagellar FlgE monomers. In panel **B**, the helical FlgE polymers are shown (zoom-in of figure A). The tiny surface pores between the flgE monomers can be seen in panels **D, E** and **F**. Panel **C** shows the view from the flagellar extracellular end and **G** shows a pore between FlgE monomers.

**Figure 7.**
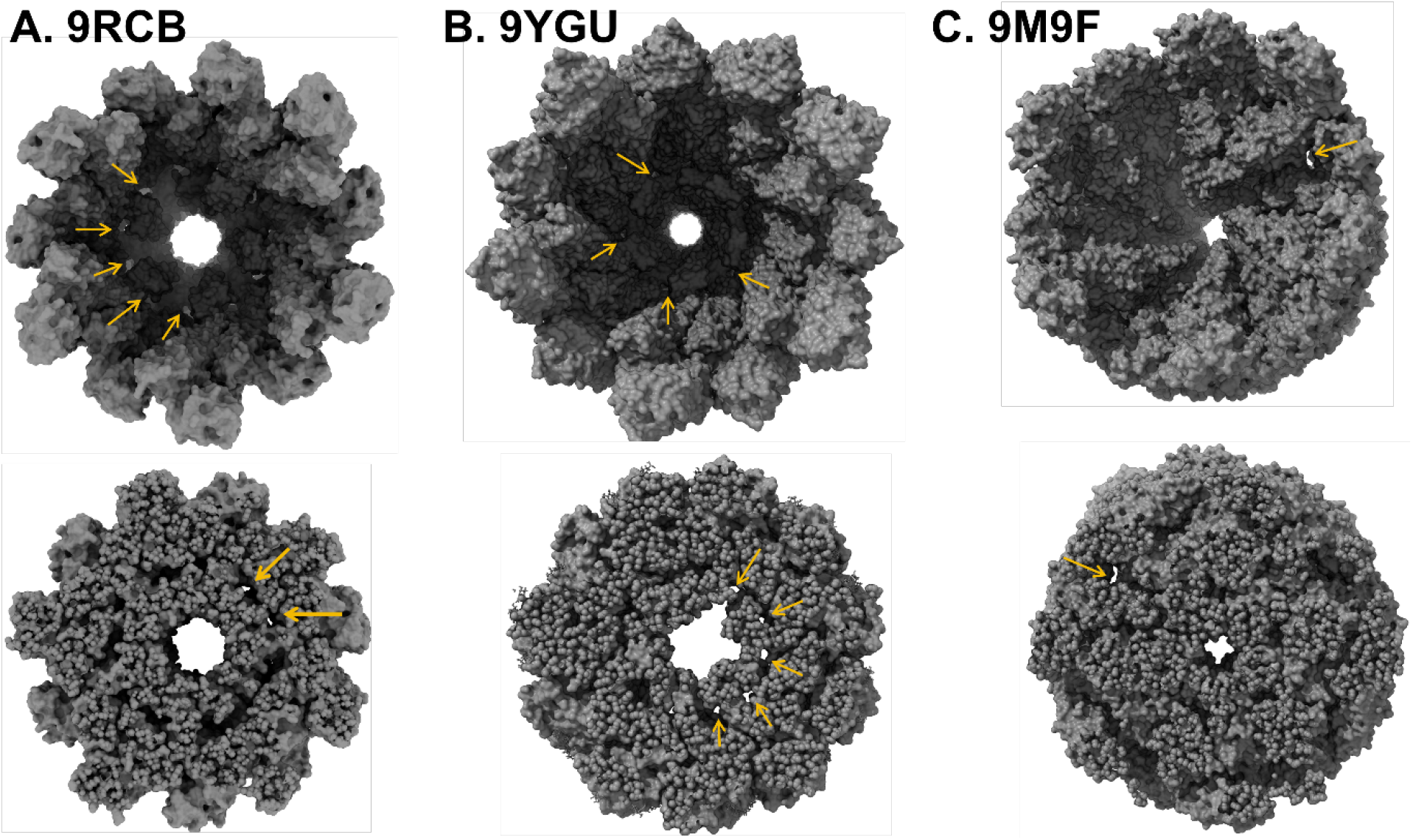
End-on and cross-sectional views of bacterial flagellar axial structures. **A –** Vibrio alginolyticus flagellar filament (PDB ID: 9RCB; Qin et al., 2026), **B**. Helicobacter pylori flagellar filament (PDB ID: 9YGU; Kumar et al., 2026) and **C**. a flagellar hook (PDB ID: 9M9F; Chen et al., 2025). Each structure presents radial canal-like inter-subunit pores surrounding the central channel; these pores measure ~10 Å in diameter, as measured using the ChimeraX distance-measurement tool. The top panels show the end-on view looking down the central channel; the bottom panels depict cross-sections of the same structures. Small radial canals can be seen (marked by yellow arrows) across all structures compared herein.

**Figure 8.**
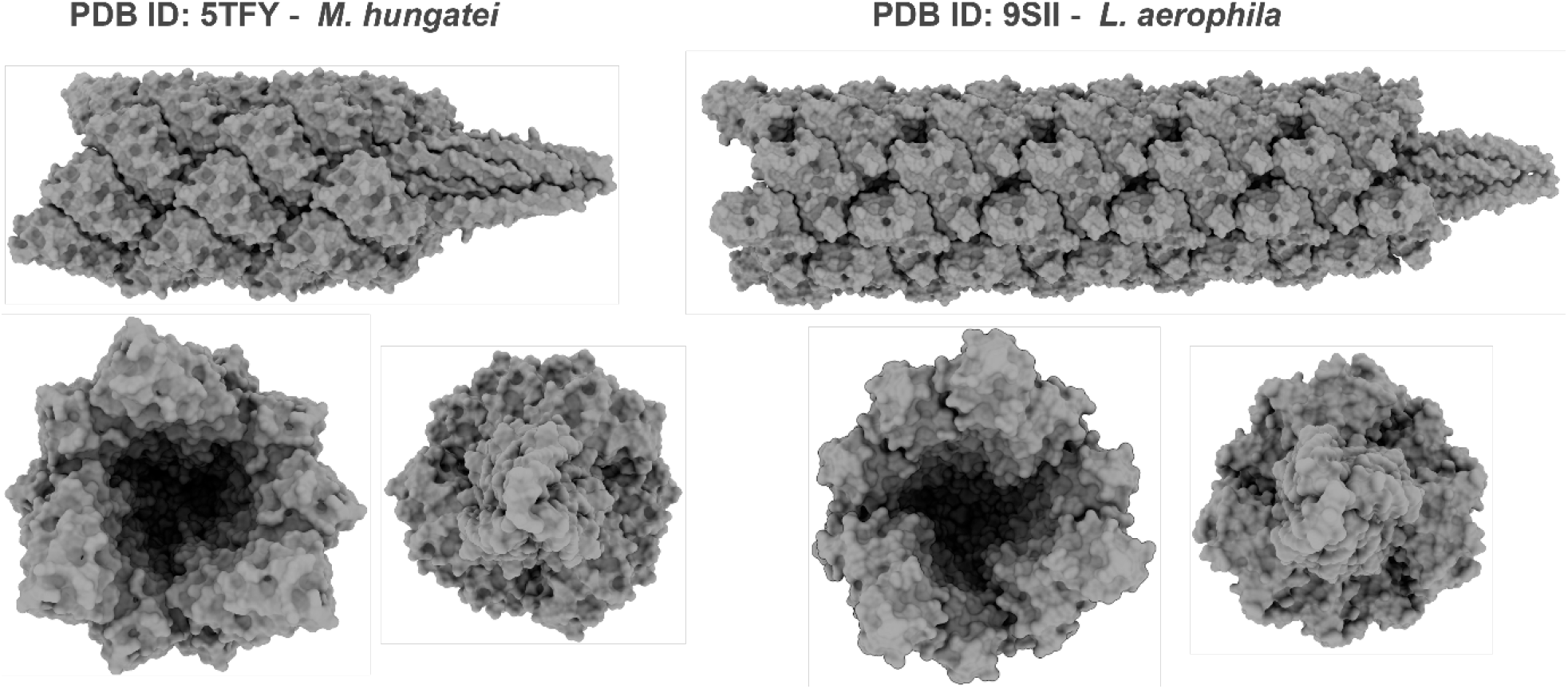
Archaellar hook structures of two archaebacterial species. **A –** M. hungatei (PDB ID: 5TFY) and **B**. L. aerophile (9SII). The top panel shows the entire hook structure. The bottom views in both panels show the view from the outer side (flagellar end) and a zoomed-in view of the pointed end seen in the top panels.

##### 3. The “Clutch/Gearbox” analogy is an anthropomorphic overreach

Analogy between the BFM and a manual transmission system in a car is usually made:

> Rotor = “molecular engine”
>
> Hook = “drive shaft” and “clutch”
>
> Stiffness change = “gear shift”

A car’s transmission is an engineered system designed by intelligence. Biological systems evolve by natural selection, not intelligent design. The analogy assumes what it needs to prove: that the system is a rotary machine. This is circular reasoning. The claimed “neutral interval” during gear shift (0.2 s) is far longer than the sub-10 ms biochemical switch – this discrepancy is hand-waved, not explained.

### IV.2 Unification of diverse motility phenomena

The murburn model proposes that all forms of microbial motility (swimming, gliding, swarming, swashing, and spirochete undulation) arise from a single physical principle: the directed ejection of metabolically produced fluid (water or hydrated slime) through hollow channels or surface pores, with passive appendages serving to harness viscous and elastic forces for thrust generation.

#### Flagellated swimming (monotrichous and peritrichous)

In monotrichous bacteria (*Vibrio cholerae*), a single polar flagellum receives water ejected from the basal T3SS module. The water exits preferentially at the hook-rod junction, creating a local shear field that induces a traveling wave along the left-handed helical filament. This wave propagates from base to tip, generating thrust via anisotropic viscous drag. The filament does not rotate axially; its apparent precession is an optical consequence of the traveling wave in shear flow, as demonstrated by Ali et al. (2016).

In peritrichous bacteria (*Escherichia coli*), multiple flagella are distributed around the cell body. During forward swimming (run), all flagella simultaneously eject water, and the filaments are drawn together by hydrodynamic attraction into a coherent bundle that trails behind the cell. The bundle propagates a collective traveling wave, generating net forward thrust. During tumbling, water ejection ceases or becomes asynchronous; the filaments relax, the bundle disperses, and the cell reorients via rotational Brownian motion. No active motor reversal is required; tumbling is simply the absence of ejection. Chemotaxis follows directly from this: rather than switching a motor between clockwise and counter-clockwise senses, the cell biases its runs by *modulating the water-ejection rate Q* in response to attractant and repellent signalling: sustaining ejection (and hence thrust) when conditions improve and pausing it to tumble when they worsen. The chemotaxis signal-transduction network thus gates a secretory flux rather than reversing a rotor, which is why the same sensory machinery operates without any directional “gearbox.”

#### Gliding motility (Type IX Secretion System)

Bacteria such as *Flavobacterium johnsoniae* glide across surfaces without any external flagellum (McBride and Nakane, 2015). The murburn model interprets gliding as water or slime ejection through surface pores of the Type IX Secretion System (T9SS). Structural studies have revealed that the T9SS core complex (GldL, GldM, GldN, PorV, SprA, SprT) exhibits circular symmetry with GldLM stator units showing 5:2 stoichiometry; reminiscent of the flagellar MotAB complex (Deme et al., 2021; James et al., 2022). In the classical view, this symmetry is interpreted as evidence for rotation of the T9SS core. In the murburn view, the circular arrangement is a pump configuration that efficiently channels slime to the cell surface. The extruded slime hydrates, expands, and flows along the cell body, creating a shear force that propels the cell forward, a “slime jet” rather than a rotating conveyor belt. **Spirochete undulation (periplasmic flagella)** Spirochetes such as *Treponema denticola* and *Borrelia burgdorferi* possess flagella that are entirely contained within the periplasmic space, between the inner and outer membranes. These periplasmic flagella are hollow and are driven by the same T3SS basal module as external flagella. In the murburn model, water ejected from the basal module flows into the periplasmic space through the hollow flagellar channel. This internal water flow creates hydraulic pressure that drives the characteristic undulatory (corkscrew) motion of the spirochete cell body. The flagellar filaments are passive conduits and structural guides; they do not rotate. This interpretation is consistent with the observation that flagella-less spirochete mutants have markedly different cell shapes (Wolgemuth et al., 2006), indicating that the flagella serve a cytoskeletal role alongside their fluid-conducting function.

#### Swarming and swashing

Swarming motility-the rapid, coordinated movement of bacterial populations across moist surfaces-involves the same water ejection mechanism as swimming, but with the cells operating at the air-liquid interface. The ejected water lubricates the surface, reducing friction, and the flagella generate thrust as in liquid swimming. The transition from swimming to swarming involves upregulation of flagellar gene expression and increased water production.

Swashing or rippling motility (observed in *Myxococcus xanthus*) involves pulsed, intermittent movement. The murburn model interprets this as periodic water ejection synchronized across a colony via chemical signaling. The on-off pulsing of ejection, not a reversal of rotation, produces the characteristic back-and-forth motion.

#### Archaea (archaellum)

Archaeal motility (archaella) is often cited as convergent evolution of a rotary motor. However, archaella are homologous to Type IV pili, not to bacterial flagella. Their filaments are solid (no central channel), so they cannot conduct a water jet. In the murburn model, archaella may use ATP-driven conformational changes to propagate traveling waves along the solid filament (like eukaryotic flagella) or the water ejection might be at the very base of the archaellum (where it pierces the bacterial body wall). In both cases, they could generate thrust without rotation. This remains to be explored/tested experimentally.

### IV.3 Experimental tests to distinguish models

Table 9 summarizes key experimental tests that can discriminate between the classical rotary model and the murburn model.

**Table 9.** Direct comparison of predictions.

| Experimental Test | Classical Rotary Prediction | Murburn Model Prediction |
| --- | --- | --- |
| Flagellum length vs. speed | Monotonic increase or plateau | Peaked optimum at native length ($\sim 10 \mu\text{m}$) |
| Water ejection rate vs. speed | No direct relationship | Speed rises with $Q$; apparent-rotation rate $f_{\text{precess}} \propto Q$ |
| Flagellar precession frequency | Equal to rotation frequency (kHz) | Traveling wave frequency; no axial rotation |
| CW rotation in still fluid | Sustained active rotation | Brief recoil only ($< 1 \text{ ms}$), if any |
| Motility with immobilized filament | Stops (rotation required) | Continues (fluid flow only) |
| Hook mutants | Altered torque transmission | Altered water ejection/shear generation |
| Uncouplers | Disrupt PMF gradient | Disrupt DRS/water production |

The murburn framework generates specific, testable quantitative predictions that distinguish it from the classical rotary model.

#### Swimming speed as a function of flagellum length

The resistive-force-theory balance of Section III sets the swimming speed; at the native length (*L*_*f*_ ≈ 10 *µ*m) it yields *U* ≈ 56 *µ*m*/*s. Because the governing dimensionless group is the sperm number Sp = *L*_*f*_ (*ξ*_⊥_*ω*_bend_*/EI*)^1*/*4^, the speed depends *non-monotonically* on *L*_*f*_ : a filament that is too short (Sp ≲ 1) behaves rigidly and develops little anisotropic drag, while one that is too long (Sp≫≪ 10) is too floppy to propagate the bending wave efficiently. The optimum therefore lies near the native length. Table 10 illustrates this dependence.

**Table 10.** Dependence of swimming speed on flagellar length (via the sperm number).

| Flagellum Length | Predicted Swimming Speed | Sperm Number | Interpretation |
| --- | --- | --- | --- |
| $L_f < 5 \mu\text{m}$ | Very low | $\text{Sp} < 3$ | Filament too short to generate sufficient anisotropic drag |
| $L_f \approx 10 \mu\text{m}$ | Maximum ($\sim 56 \mu\text{m/s}$, per Section III) | $\text{Sp} \approx 1\text{--}10$ | Optimal coupling between elastic and viscous forces |
| $L_f > 20 \mu\text{m}$ | Reduced (below peak) | $\text{Sp} > 20$ | Filament becomes too floppy; wave propagation inefficient |

Prediction 1: Engineering bacteria with flagella of different lengths (via controlled expression of FlgE mutants) should yield a peaked curve of swimming speed versus length, with maximum at the native length. The classical rotary model, treating the flagellum as a rigid propeller, would predict monotonic increase in speed with length (or no decrease beyond optimal).

#### Swimming speed and rotation rate as functions of water ejection rate

Increasing the metabolic water output *Q* raises the local shear 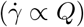 and hence the precession (apparent-rotation) rate, *f*_precess_ ∝*Q*. The swimming speed also increases monotonically with *Q*, but *not* as a simple *U*∝*Q*^2^ law: such a scaling would require a wave amplitude growing with *Q*, whereas the amplitude is in fact set geometrically and saturates (*A* ≈ *p* sin *ψ/*2*π*, Section III.6). The precise *U* –*Q* exponent therefore follows from the forced beam dynamics; the robust, model-level statements are that motility depends on metabolic water output at all (unlike a pmf-driven motor) and that the apparent-rotation rate scales linearly, *f*_precess_∝ *Q*.

Prediction 2: Manipulating metabolic water production (by varying substrate availability, oxygen tension, or inhibiting specific redox reactions) should raise the apparent-rotation rate in proportion to water output (*f*_precess_ ∝ *Q*) and monotonically increase swimming speed, a dependence already evident in the salt-dependence of the *Vibrio* sodium-driven rotation rate (Magariyama et al., 1994). Direct measurement of water ejection (e.g., using deuterated water tracers or microfluidic sensors) could test this prediction.

The fact that positive chemotaxis with usable nutrients give higher and continuous motility directly supports the murburn model (without invoking the intelligent switching of rotations!). Also, higher metabolism would give greater DROS dynamics (increasing murburn phosphorylations, thereby explaining some CheY associated observations).

#### Precession frequency and hook stiffness

Following the geometric coupling derived in Section III, the precession of the bending plane is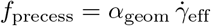, with the coupling constant *α*_geom_ ≈ 0.0056 fixed by the spiral groove pitch. For the canonical water-ejection rate *Q* ≈ 2 × 10^−20^ m^3^*/*s this gives *f*_precess_ ≈ 1,800–3,600 Hz (Section III), the rate at which the bending plane precesses about the filament axis. (Note this is distinct from the much slower bending frequency *f*_bend_ ≈78 Hz that sets the mechanical power.) The hook s effective stiffness is not an active mechanical switch but a passive consequence of the viscoelastic response to this forcing.

Prediction 3: Direct high-speed imaging of fluorescently labeled flagella should reveal traveling wave frequencies in the kHz range, not rigid body rotation at the same frequency. The wave should propagate from base to tip during forward swimming.

#### Absence of sustained CW rotation

The murburn model predicts that sustained clockwise (CW) rotation of the flagellum does not occur. What is interpreted as CW rotation in classical experiments is either: (a) a brief elastic recoil, ranging up to a few seconds, when water ejection ceases, or (b) an optical illusion from reversed shear flow direction.

Prediction 4: In experiments with truly still fluid (no convection, no thermal gradients, no external flow), the flagellum should exhibit only base-to-tip traveling waves (apparent CCW precession) during active swimming. Any CW motion should be extremely brief and non-sustained.

#### Motility without flagellar rotation

The murburn model predicts that directed swimming requires water ejection and a flexible filament, but not flagellar rotation.

Prediction 5: If the flagellar filament is physically prevented from rotating (e.g., by crosslinking agents that lock the filament in place while still allowing fluid flow through the hollow core), swimming should continue. This would directly falsify the necessity of axial rotation.

### IV.4 The hook as a flexible secretion guide: A reinterpretation

In the murburn model, the hook (FlgE polymer) is **not** a torque-transmitting universal joint. Instead, it is: (i) a flexible channel that guides water and flagellin monomers from the rod to the filament; (ii) a bending adapter that allows the filament to orient at an angle relative to the cell body (important for peritrichous bacteria); and (iii) a hydrodynamic coupler that transmits the shear flow from the basal ejection to the filament.

The variability in hook dimensions across species is **expected** in the murburn model because hook length (50-80 nm) is optimized for the specific secretion rate and filament stiffness of each species, hook diameter (14-18 nm) matches the rod diameter (14 nm) and filament core (2 nm) as a tapered channel, FlgE monomer length reflects evolutionary tuning of flexibility and channel diameter, and supercoiling state (L vs. R) determines the direction of shear-induced precession. Das et al. (2026) treat hook stiffness as a binary switch (soft for CCW, stiff for CW). In the murburn model, hook stiffness is continuously variable depending on the rate of water ejection. An elaborate critique is presented in the Supplementary Information.) It is now clear that higher Q gives higher shear, which affords higher precession frequency; and in this case the hook appears “stiffer” because it is being driven at higher frequency (viscoelastic stiffening). No active “gear shift” is required; the stiffness is an emergent property of the fluid-structure interaction.

### IV.5 Evolutionary parsimony

The murburn model offers a parsimonious evolutionary path for the origin and diversification of bacterial motility:

A. **Ancestral T3SS:** A simple secretion system evolved to export proteins across the cell envelope.
B. **Water secretion:** Metabolic redox reactions, localized near the secretion system, produced water as a byproduct. This water was co-secreted.
C. **Incidental thrust:** The ejected water created a net force on the cell, small but beneficial under certain conditions.
D. **Filament elongation:** Extensions of the secretion channel (proto-flagella) directed the water flow, increasing thrust efficiency.
E. **Helical structure:** Spontaneous polymerization of secreted proteins into helical filaments (due to their intrinsic asymmetry) created anisotropic drag, further enhancing thrust.
F. **Diversification:** Gene duplication and divergence led to different filament lengths, hook variants, and regulatory systems adapted to different environments (viscous media, surfaces, host tissues).

This evolutionary trajectory requires no “irreducibly complex” leap. Each step confers a selective advantage, however small. The classical rotary model, by contrast, requires the simultaneous emergence of a rotor, stator, ion channel, torque transmission system, and directional switch, which is a far less probable scenario.

#### The murburn model structure-function explanation

##### 1. The hook as a flexible secretion guide

In the murburn model, the hook (FlgE polymer) is unlikely to function as a torque-transmitting universal joint. Instead, it is:

> A flexible channel that guides water and flagellin monomers from the rod to the filament.
>
> A bending adapter that allows the filament to orient at an angle relative to the cell body (important for peritrichous bacteria).
>
> A hydrodynamic coupler that transmits the shear flow from the basal ejection to the filament.le

##### 2. Statistical spread reflects secretory adaptation

Variabilities in the structure-functions aspects of the BFS hook structures such as hook length (50-80 nm), hook diameter (14-18 nm) and FlgE monomer length as well as supercoiling state are interpreted differently through the murburn purview (Tables 8 and 11).

**Table 11.** Classical vs. murburn model of hook morphology.

| Prediction | Classical Rotary Model | Murburn Model |
| --- | --- | --- |
| Hook stiffness should be conserved across species | Yes (must be precise for torque transmission) | No (varies with secretion rate and filament length) |
| Hook length should correlate with torque output | Yes | No; correlates with filament length and water ejection rate |
| Hook mutants with altered flexibility should show changed torque-speed curves | Yes | Yes, but the mechanism is altered shear coupling, not torque transmission |
| Water flow through hook should be detectable | No (hook only transmits torque) | Yes (hook is a fluid channel) |

##### 3. Hook stiffness-not a binary switch

Das et al. treat hook stiffness as a binary switch (soft for CCW, stiff for CW). In the murburn model:

Hook stiffness is continuously variable depending on the rate of water ejection.

Higher Q→ higher shear→ higher precession frequency→ the hook appears “stiffer” because it is being driven at higher frequency (viscoelastic stiffening).

No active “gear shift” is required – the stiffness is an emergent property of the fluid-structure interaction.

##### 4. The “universal joint” analogy is replaced by “flexible hose”

The classical “universal joint” analogy (hook as a mechanical coupler) is replaced by the **flexible hose** analogy:

> A flexible hose transmits fluid from a pump to a nozzle. The hose can bend without breaking, and the fluid flow does not require the hose to rotate.
>
> The flagellar hook is a **biological hose** – it channels water from the basal T3SS to the filament nozzle. Its flexibility allows the filament to orient at various angles without kinking.

##### 5. Testable predictions for hook morphology (Murburn vs. Classical)

When comparing aspects such as the stiffness of the hook, correlation between length & torque output, as well as fluid transport within the channel, there are remarkable differences as highlighted in Table 11.

In the overview, the Das et al. (2026) paper, while mathematically sophisticated, does not pose merit in that it fails to discuss earlier murburn-based criticisms and is built on the faulty foundation of the rotary motor paradigm. Their treatment of the hook as a mechanical universal joint and torque diverter (Table 12) is an overinterpretation that does not account for the statistical spread of hook morphology across bacterial species.

**Table 12.** Classical vs. murburn interpretation of the hook.

| Aspect | Das et al. (Classical) | Murburn Model |
| --- | --- | --- |
| Primary function | Torque transmission (universal joint) | Fluid channel (secretion guide) |
| Stiffness regulation | Active mechanical switch (CCW soft, CW stiff) | Passive viscoelastic response to shear rate |
| Bending angle $\psi$ | Determined by torque balance | Determined by water ejection rate and filament drag |
| Statistical spread | Ignored or treated as noise | Expected; reflects evolutionary tuning of secretion |
| Energy source | PMF (proton gradient) | Metabolic water production (redox) |
| Directionality | CCW vs. CW active rotation | CCW (base→tip wave) active; CW (recoil) passive |
| Gearbox analogy | Central to model | Anthropomorphic overreach |

In the murburn model:

> The hook is a flexible secretion channel, not a torque transmitter.
>
> Its stiffness is a passive emergent property of water ejection rate and filament dynamics, not an active mechanical switch.
>
> The variability in hook dimensions across species is expected and reflects evolutionary tuning to different secretion rates, filament lengths, and environmental conditions.

We propose that the hook should be understood as a biological flexible hose – a simple, evolvable, and parsimonious structure that channels water from the basal pump to the filament nozzle, enabling motility without any rotary components.

## V. Conclusions

The quantitative backbone of the argument is a single force-balance comparison of the three candidate mechanisms (Section III.9): of direct jetting, rigid helical rotation, and shear-driven elastohydrodynamics, only the last meets the ≈0.8 pN thrust requirement with metabolically sustainable water and without a rotary motor. Direct jetting fails by ~6 orders of magnitude on thrust, and the water flux it would need drains the cell in ~0.02 s, while rigid rotation demands precisely the proton-motive engine that the preceding sections show to be untenable.

Quantitatively, in a unipolar flagellated model the metabolically sustained water ejection (*Q* ≈ 2 × 10^−20^ m^3^*/*s, *v*_ej_ ≈ 6.4 × 10^−3^ m*/*s) generates a local shear 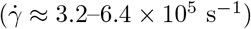 that the filament’s spiral grooving couples into a transverse bending wave (bending frequency *f*_bend_ ≈ 78 Hz; precession of the bending plane *f*_precess_ ≈ 1,800–3,600 Hz). Resistive-force theory then yields a thrust *F*_thrust_ ≈0.8 pN and a swimming speed *U* ≈30–56 *µ*m*/*s, within the observed range, at a mechanical cost of ≈0.15 pW per flagellum, a small fraction (*<* 5%) of the cellular metabolic budget.

The classical rotary model of bacterial flagellar motility, despite decades of acceptance, faces insurmountable challenges:

1. Thermodynamic impossibility: The proton budget exceeds cellular capacity by 10^5^–10^6^-fold. At physiological pH, a bacterium contains *<*10 free protons, yet the classical model requires ~10^6^ protons/second.
2. Structural implausibility: A multi-layer rotating shaft with sub-Angstrom seals cannot exist in a fluid, thermally agitated membrane. Stator rotation at 13,600 Hz (816,000 RPM) in a viscous lipid bilayer is mechanically absurd.
3. Lack of sequence conservation: Purported “essential” rotary residues (e.g., D32 in MotB) are not conserved across diverse bacterial species (Gideon et al., 2024, Table 4). What is conserved is a high proportion of charged/polar residues, consistent with DRS recruitment, not rotation.
4. Kinematic inconsistency: The left-handed screw geometry predicts the opposite thrust direction from observations. The classical model s reliance on low-Re hydrodynamics to “resolve” this is an admission that the screw analogy fails.
5. Evidence-reinterpretation: Gold bead “rotation” can be explained by passive hydrodynamic effects in shear flow (Ali et al., 2016). No direct visualization of axial rotation exists at optical resolution (20 nm filament vs. 400–700 nm light).
6. Evolutionary parsimony: The flagellar basal body is homologous to the T3SS, which does not rotate. Parsimony dictates that neither rotates; both are secretion systems. The classical model requires distinct mechanisms for flagellated swimmers, gliders, spirochetes, and archaea. The murburn model explains all with one principle.

The murburn model offers a parsimonious, physically defensible, and experimentally testable alternative:

- Water is produced locally in the C-ring via DRS-mediated redox reactions (the “water-producing crucible”).
- Local heating reduces viscosity and creates pressure.
- Ejection through hollow filament generates thrust via fluid displacement at low Re (this ejection would also be needed for flagellin to move through to the tip of the filament!)
- No rotation is required; apparent rotation is a passive hydrodynamic effect of helical filaments in flow.
- All motility types (swimming, gliding, spirochete undulation, archaella) are unified under this single mechanism.

The murburn model does not merely challenge the rotary paradigm; it replaces it with a simpler, more coherent framework that aligns with thermodynamics, structural biology, fluid mechanics, evolutionary reasoning, and direct observational constraints.

The murburn model does not yet provide a fully quantitative description of peritrichous bundling or the transition from swimming to swarming, does not pinpoint the exact loci of water ejection and does not address the details of archaeal motility. These are areas for future work. The model would be falsified by: (i) direct visualization of sustained axial rotation of the flagellum in still fluid; (ii) demonstration that swimming continues when the hollow core of the filament is blocked; (iii) measurement of water ejection rates that are inconsistent with metabolic water production.

## Appendix 1: Derivation of key equations

### A1. Shear rate from basal flow

For a nozzle of radius *r*_*n*_, volumetric flow *Q*, the velocity at the nozzle exit is 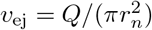. The velocity gradient at the filament surface (distance *r*_*f*_ from the centerline) is approximated by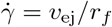 for *r*_*f*_ ≫ *r*_*n*_. If the flow is fully developed, the exact shear rate at the wall ofa pipe of radius *r*_*f*_ is 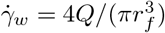 */*(*πr*^3^). For *r*_*f*_ = 10 nm, this gives 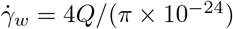. For *Q* = 4 × 10^−20^,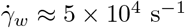– still sufficient.

### A2. Resistive-force theory derivation

For a slender filament, the local drag anisotropy yields the net swimming speed. The full derivation (Lighthill, 1976) gives:

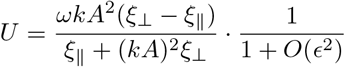

where *ϵ* = *r*_*f*_ */λ* is the slenderness parameter. For bacterial flagella, *ϵ* ≪≪1, so the expression is accurate to within a few percent.

### A3. Sperm number and optimal propulsion

The sperm number arises from the balance of viscous and elastic forces. The optimal range *Sp*~ 1− 10 ensures that the wave amplitude is not damped out (too high frequency) nor too slow to generate thrust (too low frequency). This is analogous to the condition for maximum swimming efficiency in eukaryotic flagella and cilia.

## Appendix 2: We shall critique a recent manuscript, which advocates the pmf-based operability of rotary flagella, Das et al. (2026)

### The bacterial flagellar hook as a PMF-driven motor vs. murburn view

#### 1. Fundamental premise is untenable

The entire model rests on the assumption that the bacterial flagellar motor (BFM) is a **rotary engine** driven by **proton motive force (PMF)**. In contrast, the murburn model questions the classical assumptions and presents logical arguments to rethink the structure-function aspects of the BFS (Table 8). As we have extensively demonstrated in our previous works (Gideon et al., 2024; Manoj and Bazhin, 2021; Manoj et al., 2023a,b,c,d), the PMF-based bioenergetic framework is thermodynamically, kinetically, and structurally untenable. Tables 11 and 12 present the classical vs. murburn view of the bacterial hook.

## Appendix 3: Kinematic visualisation script for the murburn model

### What this script is, and is not

The MATLAB script below renders the closed-form relations of Section III as a moving picture: the cell body is translated at the analytically predicted swimming speed *U* while the filament is drawn carrying the geometrically derived wave (amplitude *A* = *p* sin *ψ/*2*π*, wave count *n*_waves_). It is provided purely as a *kinematic aid to intuition*. It is **not** a fluid-dynamics simulation: it does not solve the Stokes equations, computes no flow field, and performs no force balance beyond the algebra already given in the main text. It therefore constitutes **neither a validation nor a proof** of the model; its only role is to make the predicted motion visible. Three points make the display-only nature explicit and reproducible:

### Frequencies are slowed for the eye

The physical precession is in the kHz range, far too fast to perceive. The animation multiplies it by a fixed display factor (≈3 × 10^−3^) so the wave can be seen; the on-screen oscillation rate is thus a viewing convenience, *not* the physical frequency.

### The parameter block reproduces Section III

The first part of the script recomputes *v*_ej_, 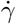, *α*_geom_, *f*_precess_, *f*_bend_ and *A* from the same inputs used in the text and prints them. For the swimming speed it prints both ends of the Section III.6 predicted range (≈30–56 *µ*m/s): the first-principles force-free self-propulsion value (lower) and the thrust-vs-body-drag value (upper, used to drive the animation). This is an internal arithmetic check, not an independent test of the physics.

### The flow-rate sweep illustrates speed saturation

Because the wave amplitude is geometric, the swimming speed is essentially *Q*-independent (Section III.6); the *Q*-sweep therefore holds *U* fixed while the precession rate *f*_precess_ scales with *Q*, visualising the saturation prediction rather than any *U* (*Q*) law.

### The on-screen wave is a planar simplification

The animation renders the filament as a planar travelling wave; the model’s actual kinematics are a *precessing* bending plane (the “apparent rotation”), so the drawing conveys the travelling-wave motion but not the full helical precession geometry.

No machine-specific paths or hidden inputs are used: every parameter is the value stated in Section III, and output is written to a folder created in the current working directory, so the script runs unmodified on any platform.

**Figure L1.**
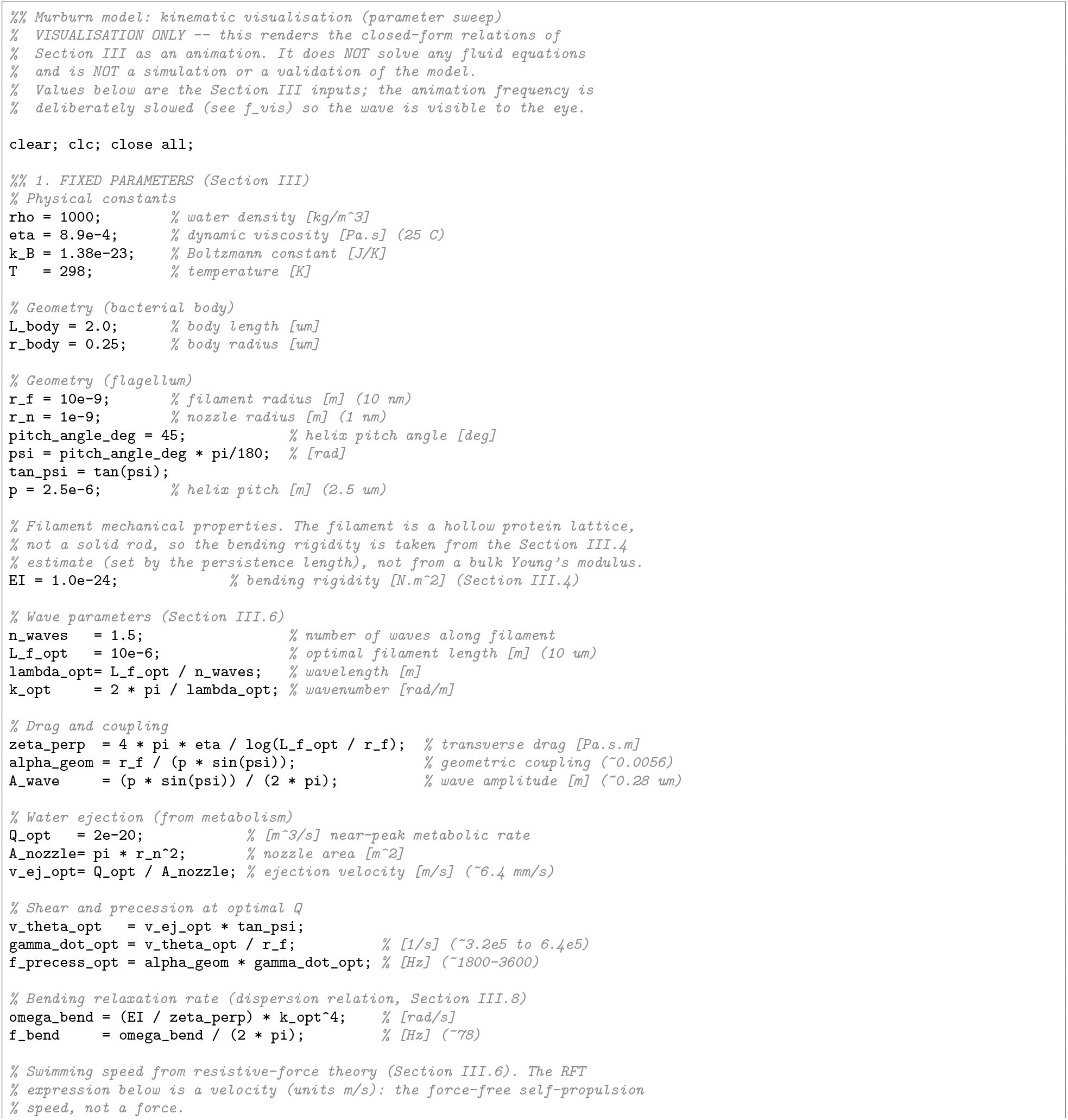
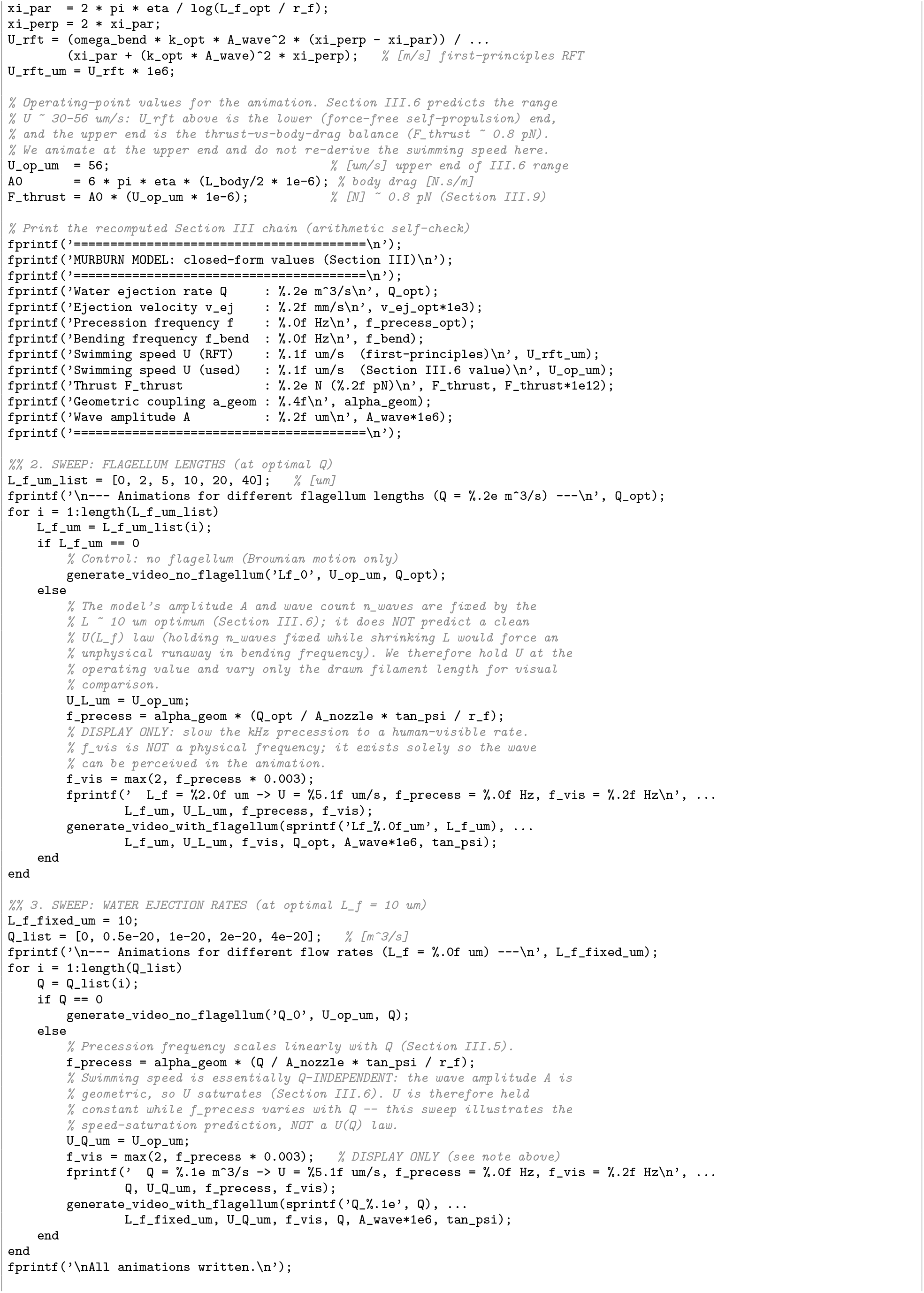
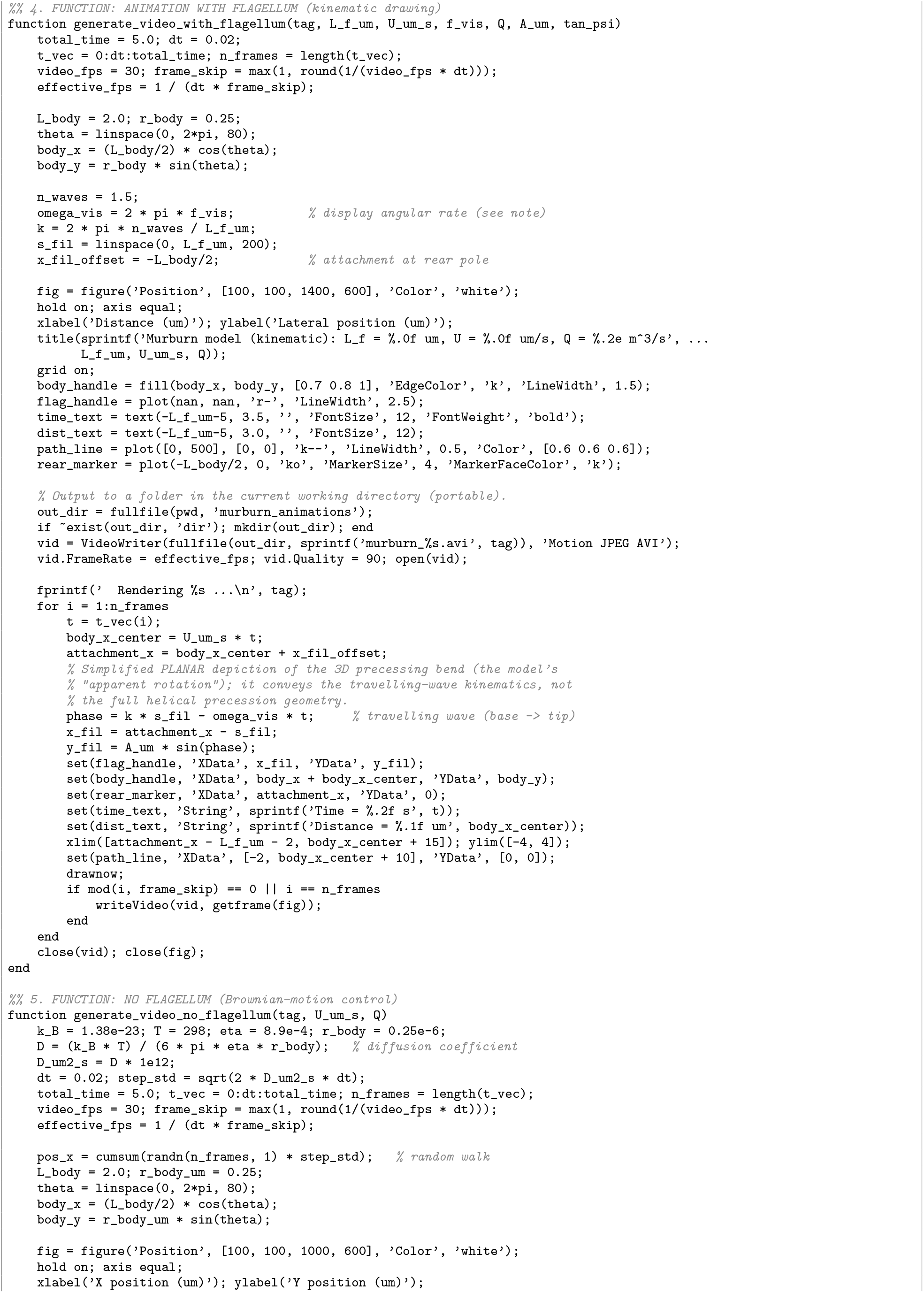
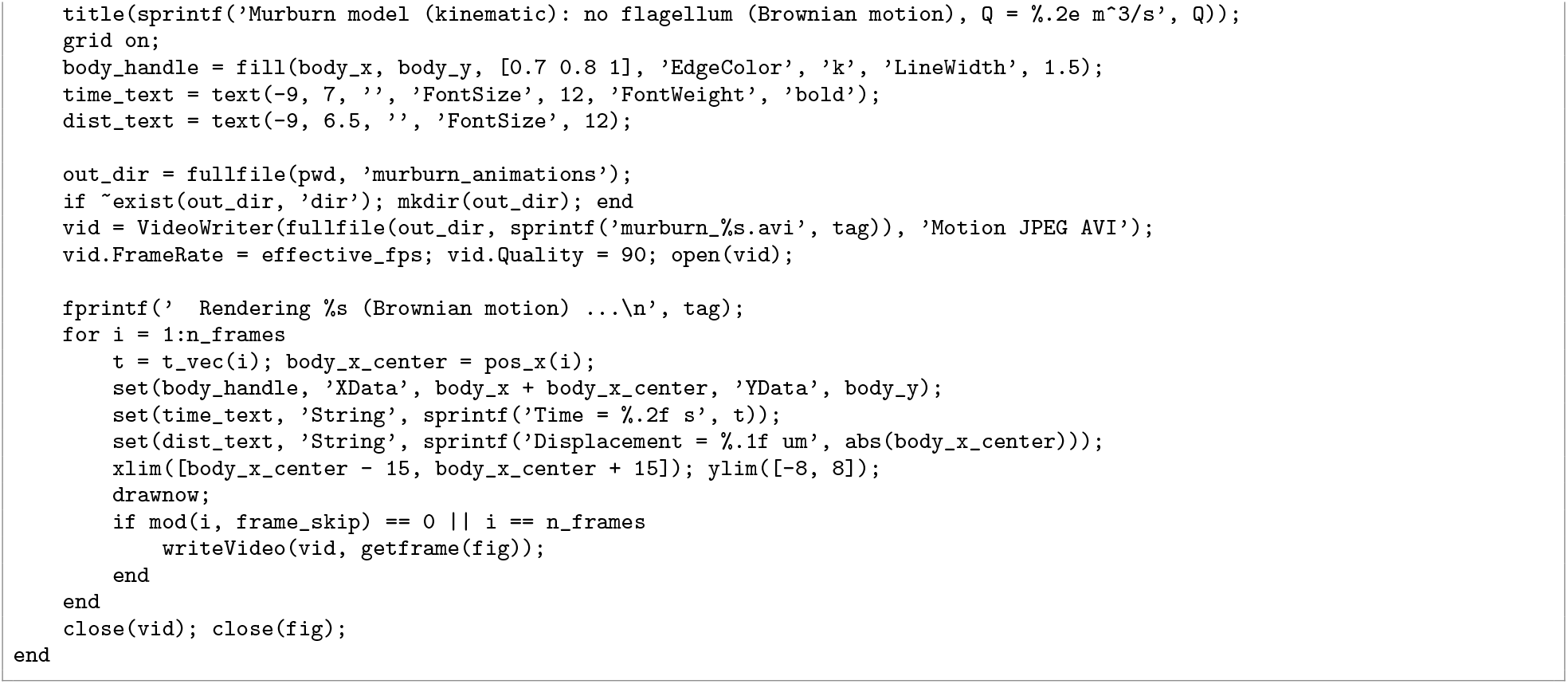
Kinematic visualisation of the murburn model (display-only; not a hydrodynamic simulation).

